# Statistics of nascent and mature RNA fluctuations in a stochastic model of transcriptional initiation, elongation, pausing, and termination

**DOI:** 10.1101/2020.05.13.092650

**Authors:** Tatiana Filatova, Nikola Popovic, Ramon Grima

## Abstract

Recent advances in fluorescence microscopy have made it possible to measure the fluctuations of nascent (actively transcribed) RNA. These closely reflect transcription kinetics, as opposed to conventional measurements of mature (cellular) RNA, whose kinetics is affected by additional processes downstream of transcription. Here, we formulate a stochastic model which describes promoter switching, initiation, elongation, premature detachment, pausing, and termination while being analytically tractable. By computational binning of the gene into smaller segments, we derive exact closed-form expressions for the mean and variance of nascent RNA fluctuations in each of these segments, as well as for the total nascent RNA on a gene. We also derive exact expressions for the first two moments of mature RNA fluctuations, and approximate distributions for total numbers of nascent and mature RNA. Our results, which are verified by stochastic simulation, uncover the explicit dependence of the statistics of both types of RNA on transcriptional parameters and potentially provide a means to estimate parameter values from experimental data.

## 1 Introduction

Transcription, the production of RNA from a gene, is an inherently stochastic process. Specifically, the interval of time between two successive transcription events is a random variable whose statistics depend on multiple single-molecule events behind transcription [1]. When the distribution of this random variable is exponential, we say that expression is constitutive; in that case, the number of transcripts produced in a certain interval of time follows a Poisson distribution. On the other hand, when the distribution of times between two successive transcripts is non-exponential, then the number of transcripts is non-Poissonian. A special case of such non-constitutive behaviour is bursty expression, whereby transcripts are produced in short bursts that are separated by long silent intervals [2,3]. In yeast, genes whose expression is constitutive include MDN1, KAP104, and DOA1, whereas PDR5 is an example of a gene whose expression is bursty [4].

For two decades, mathematical models of gene expression have been developed to predict the distribution of RNA abundance. By matching the theoretical distribution with experimental measurements from microscopy-based methods [5], one hopes to obtain insight into the underlying kinetics of transcription, and to estimate transcriptional parameters. The standard model of gene expression which has been used for these analyses is the telegraph model [6], whereby a gene can be in two states. Transcription occurs in one of the states, whereupon RNA degrades; first-order kinetics is assumed for all processes. While the distribution obtained from the telegraph model can typically fit cellular RNA abundance data, there are innate difficulties with the interpretation of that fit: fluctuations in cellular RNA numbers and, hence, the shape of the experimental RNA distribution do not only reflect transcription, but also many processes downstream thereof, such as splicing, RNA degradation, and partitioning during cell division.

To counteract these difficulties, in the past few years, mathematical models [7–10] have been developed to predict the statistics of nascent RNA, i.e. of RNA in the process of being synthesized by the RNA polymerase molecule (RNAP), which can be visualized and quantified due to recent advances in fluorescence microscopy [11–15]. In contrast to cellular RNA, the statistics of nascent RNA is a direct reflection of the transcription process; hence, these models can potentially give more insight than the simpler, but cruder telegraph model. Choubey and collaborators [7,8] have developed a stochastic model with the following properties: (i) a gene can be in two states (active or inactive); (ii) from the active state, transcription initiation occurs in two sequential steps: the pre-initiation complex is formed, after which the RNA polymerase escapes the promoter; (iii) once on the gene, the polymerase moves from one base pair to the next (with some probability) until the end of the gene is reached, when transcription is terminated and polymerase detaches. Queuing theory is used to derive analytical expressions for the transient and steady-state mean and variance of numbers of RNAP that are attached to the gene in the long-gene limit when the elongation time is practically deterministic. Xu et al. [9] have considered a coarse-grained version of that model, whereby the movement of RNAP from one base pair to the next is not explicitly modeled, obtaining an analytical expression for the total RNAP distribution in steady-state conditions. More recently, Cao and Grima [10] have studied a model of eukaryotic gene expression that yields approximate time-dependent distributions of both nascent and cellular RNA abundance as a function of the parameters controlling gene switching, DNA duplication, partitioning at cell division, gene dosage compensation, and RNA degradation; in their coarse-grained model, the movement of RNAP is not explicitly modeled, while the elongation time is assumed to be exponentially distributed, which simplifies the requisite analysis.

The complexity of nascent RNA models has thus far not allowed the same detailed level of analysis as has been possible with the much simpler telegraph model. A few shortcomings of current models can be summarized as follows: (i) distributions of nascent RNA have been derived from models that do not explicitly model the movement of RNAP along a gene [9,10], resulting in a disconnect between theoretical description and the microscopic processes underlying transcription; (ii) while the analysis of single-cell sequencing data and electron micrograph data yields the positions of individual polymerases along the gene, allowing for the calculation of statistics (means and variances) of the numbers of RNAP on gene segments that are obtained after binning, detailed models of RNAP elongation [7,8] provide analytical results only for total RNAP on a gene and hence cannot be used to understand gene segment data; (iii) analytical calculations of the statistics of nascent RNA ignore important details of the transcription process such as pausing, traffic jams, backtracking, and premature termination, some of which have to-date been explored via stochastic simulation [7,16–19].

In this paper, we overcome some of the aforementioned shortcomings of analytically tractable models for transcription. In Section 2, we study a stochastic model for promoter switching and the stochastic movement of RNAP along a gene, allowing for premature termination. We derive exact closed-form expressions for the first and second moments (means and variances) of local RNAP fluctuations on gene segments of arbitrary length, which allows us to study how these statistics vary along a gene as a function of transcriptional parameters; we also obtain expressions for the mean and variance of the total RNAP on the gene which generalize previous work by Choubey et al. [7]. In Section 3, we investigate approximations for the distributions of total RNAP and mature RNA, showing in particular that Negative Binomial distributions can provide an accurate approximation in certain biologically meaningful limits. In Section 4, we illustrate the difference between the statistics of local and total RNAP fluctuations and those of light fluorescence due to tagged nascent RNA. In Section 5, we extend our model to include pausing by deriving approximate expressions for the mean, variance, and distribution of observables. We conclude with a discussion of our results in Section 6.

## 2 Detailed stochastic model of transcription: setup and analysis

In this section, we specify the stochastic model studied here; then, we derive closed-form expressions for the moments of mature RNA and local and total RNAP fluctuations in various parameter regimes.

### 2.1 Setup of model

We consider a stochastic model of transcription that includes the processes of initiation, elongation, and termination, as illustrated in Fig. 1. For simplicity, we divide the gene into *L* segments; the RNAP on gene segment *i* is then denoted by *P*_*i*_. The promoter can be either in the inactive state (*G*_off_) or the active state (*G*_on_), switching from the inactive state to the active one with rate *s*_*u*_ and from the active state to the inactive one with rate *s*_*b*_. When promoter is active, initiation commences via the binding of an RNAP wtih rate *r*, denoted by *P*_1_. Subsequently, the RNAP either moves from a gene segment to the neighboring segment with rate *k*, or it prematurely detaches with rate *d*. Note that here we have made two assumptions: (i) the movement of RNAP is unidirectional, away from the promoter site and hence left to right, with no pausing or backtracking allowed; (ii) the detachment and elongation rates are independent of the position of RNAP on the gene. Each RNAP has associated with it a nascent RNA tail that grows longer as the RNAP transcribes more of the gene. When the RNAP reaches the last gene segment, termination occurs, i.e. the RNAP-nascent RNA complex gets dissociated from the gene leading to a mature RNA (*M*) which degrades with rate *d*_*m*_. Note that for simplicity, we have not considered excluded-volume interaction between adjacent RNAPs here; hence, we make the implicit assumption of low ‘traffic’, which is plausible when the initiation rate is sufficiently low.

**Figure 1:**
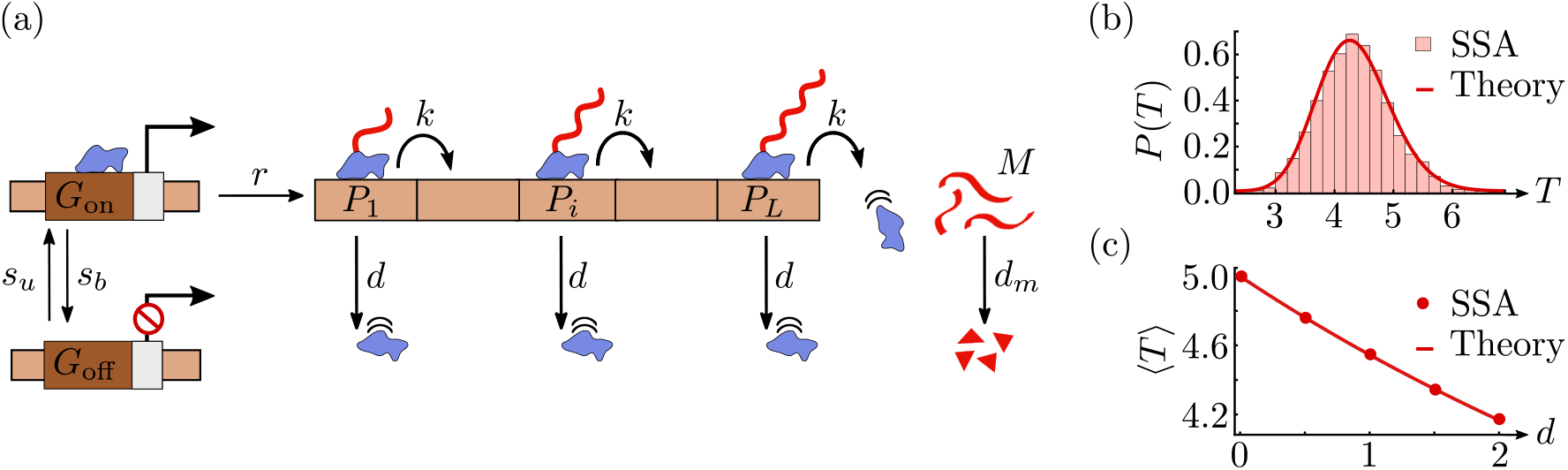
Model of transcription. (a) The gene is arbitrarily divided into *L* segments, with RNAP (blue) on gene segment *i* denoted by *P*_*i*_. The promoter switches from the active state *G*_on_ to the inactive state *G*_off_ with rate *s*_*b*_, while the reverse switching occurs with rate *s*_*u*_. When the promoter is active, initiation of RNAP occurs with rate *r*. Initiation is followed by elongation, which is modelled as RNAP ‘hopping’ from gene segment *i* to the neighbouring segment *i* + 1 with rate *k*, i.e. as the transformation of species *P*_*i*_ to *P*_*i*+1_. RNAP prematurely detaches from the gene with rate *d*. A nascent RNA tail (red), attached to the RNAP, grows as elongation proceeds. Termination is modelled by the change of *P*_*L*_ with rate *k* to mature RNA (*M*), which subsequently degrades with rate *d*_*m*_. In panel (b), we show the probability distribution *P*(*T*) of the total elongation time *T* – the time between initiation and termination – as predicted by the stochastic simulation algorithm (SSA; histogram) and our theory (Erlang distribution with shape parameter *L* and rate *k* + *d*; solid line). The parameter values used are *L* = 50, *k* = 10/min, and *d* = 1.5/min. In panel (c), we show the dependence of the mean of the distribution *P*(*T*) on the RNAP detachment rate (*d*), as predicted by SSA (dots) and our theory (〈*T*〉 = *L*/(*k* + *d*); solid line). The relevant parameter values are *L* = 50 and *k* = 10/min.

The elongation time which is the total time *T* from initiation to termination, that is, conditioning on those realisations for which the RNAP does not prematurely detach, is Erlang distributed with mean *L*/(*k* + *d*) and coefficient of variation 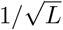; see Appendix A for a derivation and Figs. 1(b) and (c) for verification through stochastic simulation (SSA).

Note that the total number of RNAPs transcribing the gene is equal to the number of nascent RNA molecules present, *irrespective of their lengths*; to shed light on the fluctuations of nascent RNA, in this section we therefore focus on the calculation of statistics of local and total RNAP fluctuations. We define the vector of molecule numbers 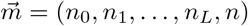, and we write 〈*n*_0_〉, 〈*n*_*i*_〉 (*i* = 1, 2, …, *L*), and 〈*n*〉 for the average numbers of molecules of active gene, RNAP, and mature RNA, respectively. The above model can then be conveniently described by *L* + 2 species interacting via a set of 2*L* + 4 reactions with the following rate functions:

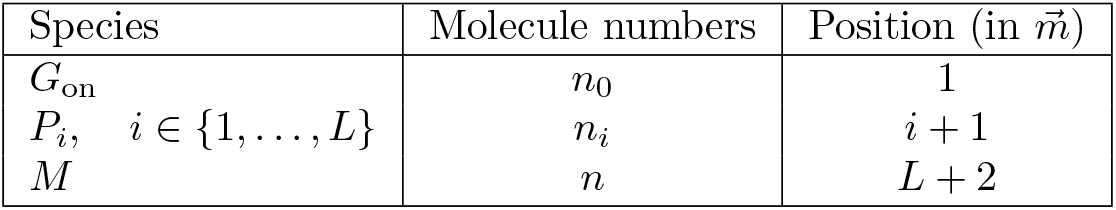

| Reaction | Rate function $f_j$ |
| --- | --- |
| $G_{\text{on}} \xrightarrow{s_b} G_{\text{off}}$ | $f_1 = s_b \langle n_0 \rangle$ |
| $G_{\text{off}} \xrightarrow{s_u} G_{\text{on}}$ | $f_2 = s_u (1 - \langle n_0 \rangle)$ |
| $G_{\text{on}} \xrightarrow{r} G_{\text{on}} + P_1$ | $f_3 = r \langle n_0 \rangle$ |
| $P_i \xrightarrow{k} P_{i+1}, \quad i \in \{1, \dots, L-1\}$ | $f_{i+3} = k \langle n_i \rangle$ |
| $P_L \xrightarrow{k} M$ | $f_{L+3} = k \langle n_L \rangle$ |
| $P_i \xrightarrow{d} \emptyset, \quad i \in \{1, \dots, L\}$ | $f_{i+L+3} = d \langle n_i \rangle$ |
| $M \xrightarrow{d_m} \emptyset$ | $f_{2L+4} = d_m \langle n \rangle$ |

Note that *G*_off_ is not an independent species; the reason is that the binary state of the gene implies a conservation law, with the sum of the numbers of *G*_on_ and *G*_off_ equaling 1. Hence, the number of independent species in the model is *L* + 2. The rate functions *f*_*j*_ are the averaged propensities from the underlying chemical master equation (CME); note that, because our reaction network is composed of first-order reactions, these rate functions also equal the reaction rates in the corresponding deterministic rate equations. The description of our model is completed by the (*L*+2)×(2*L*+4)-dimensional stoichiometric matrix **S**; the element **S**_*ij*_ of **S** gives the net change in the number of molecules of the *i*-th species when the *j*-th reaction occurs. Given the ordering of species and reactions as described in the Tables above, it follows that the matrix **S** has the simple form

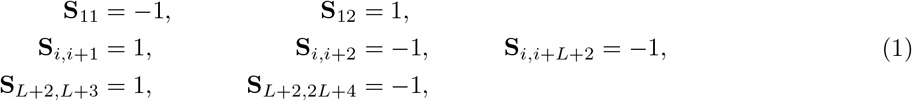

where *i* = 2, …, *L* + 1.

### 2.2 Closed-form expressions for moments of mature RNA and local RNAP

In this subsection, we outline the derivation of the steady-state means and variances of local RNAP fluctuations (on each gene segment), as well as of mature RNA. Our results are summarized in the following two propositions.

#### Proposition 1.

*Let η* = *s*_*u*_/(*s*_*u*_ + *s*_*b*_) *be the fraction of time the gene spends in the active state, let ρ_k_* = *r/k be the mean number of RNAPs binding to the promoter site in the time it takes for a single RNAP to move from one gene segment to the next, let ρ* = *r/d_m_ be the mean number of RNAPs binding to the promoter site in the time it takes for a mature RNA to decay, and let μ* = *k*/(*k* + *d*) *be the probability that RNAP moves to the next gene segment rather than detaching prematurely. Then, the steady-state mean numbers of molecules of active gene, RNAP, and mature RNA are given by*

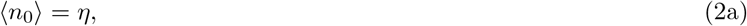

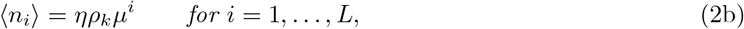

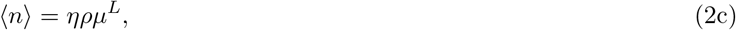

*respectively.*

Prop. 1 can be proved in a straightforward fashion, as follows. Using the underlying CME, one can show from the corresponding moment equations [20] that the time-evolution of the vector 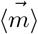 of mean molecule numbers in a system of zeroth-order or first-order reactions, i.e. with propensities that are linear in the number of molecules, is given by the time derivative 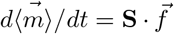. Given the form of the stoichiometric matrix **S** and of the rate functions *f*_*j*_, as described in Section 2.1, it follows that the mean numbers of all species in steady-state can be obtained by solving the following system of *L* + 2 algebraic equations:

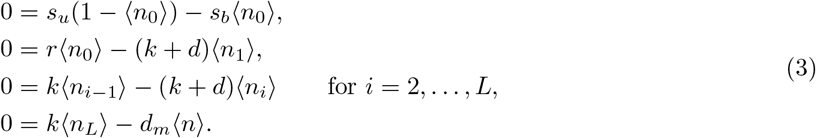

These equations can easily be solved simultaneously to yield the steady-state value of 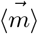, as given in Eq. (2).

#### Proposition 2.

*Let τ_p_* = 1/(*d* + *k*), *τ*_*g*_ = 1/(*s*_*u*_ + *s*_*b*_), *and τ*_*m*_ = 1/*d*_*m*_ *be the timescales of fluctuations of RNAP, gene, and mature RNA, respectively, such that we can define the three new parameters*

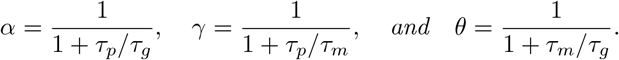

*Furthermore, let β* = *s*_*b*_/*s*_*u*_ *denote the ratio of gene inactivation and activation rates. Then, the variances and covariances of molecule number fluctuations of active gene, RNAP, and mature RNA are given by*

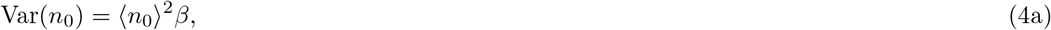

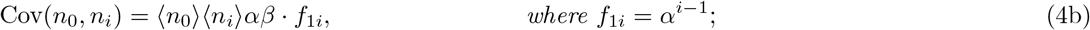

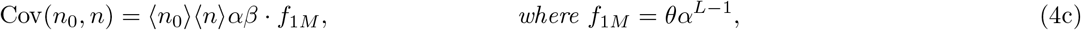

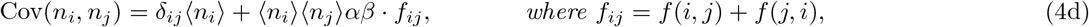

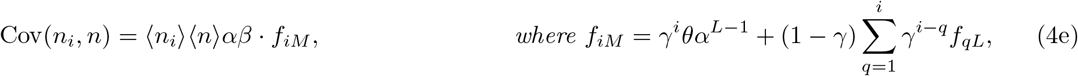

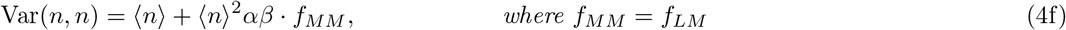

*for i, j* ∈ {1, …, *L*}. *Here, δ_ij_ is the Kronecker delta function; moreover,*

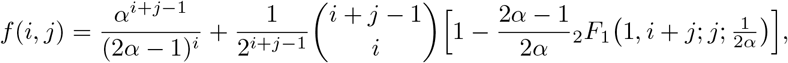

*where* _2_*F*_1_ *denotes the generalized hypergeometric function of the second kind [21], which is defined as*

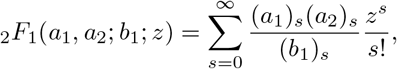

*with* (*a*)_*s*_ = Γ(*a* + *s*)/Γ(*a*) *the Pochhammer symbol.*

As above, since the underlying propensities are linear in the number of molecules, the CME implies [20] that the corresponding second moments in steady-state are exactly given by a Lyapunov equation. That equation, which is precisely the same as the one that is obtained from the linear-noise approximation (LNA) [22], takes the form

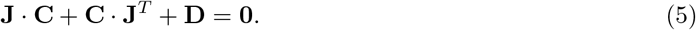

Here, **C**, **J**, and **D** are (*L* + 2) × (*L* + 2)-dimensional matrices; **C** is a variance-covariance matrix that is symmetric (**C**_*ij*_ = **C**_*ji*_), **J** is the Jacobian matrix with elements 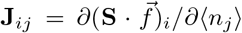, and 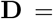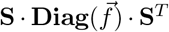 is a diffusion matrix, where 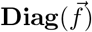 is a diagonal matrix whose elements are the entries in the rate function vector 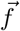. The non-zero elements of **J** are given by

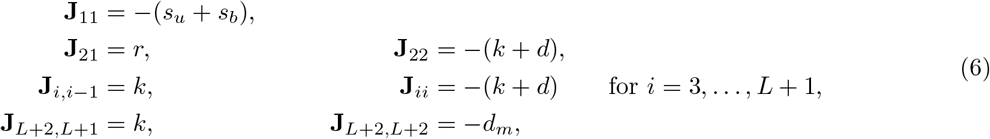

while the non-zero elements **D**_*i*_ read

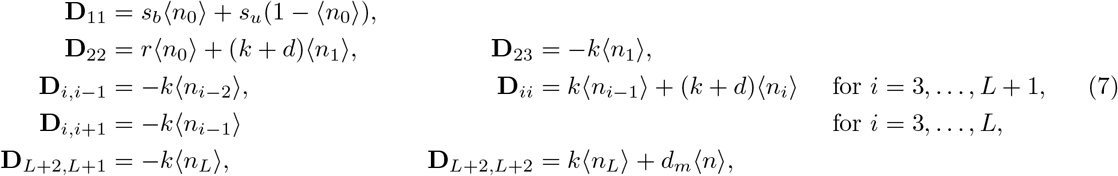

Given the structure of the matrices **J** and **D** above, the Lyapunov Eq. (5) can be solved explicitly for the covariance matrix **C** whose elements are given by Eq. (4). The solution by induction is involved and can be found in Appendix B, which proves Prop. 2.

#### 2.2.1 Simplification in bursty and constitutive limits

##### Bursty limit

We now consider a particular parameter regime – the limit of large initiation rate *r* and large gene inactivation rate *s*_*b*_ such that *b* = *r*/*s*_*b*_ is constant. Since the fraction of time spent in the active state is *η*, it follows that the gene is mostly in the inactive state in that limit. During the short periods of time when it transitions to the active state, a burst of initiation events occur; in particular, a mean number *b* of RNAPs bind to the promoter during activation. Hence, such genes are often termed bursty, since transcription proceeds via sporadic bursts of activity and *b* is called the mean transcriptional burst size. For *r* and *s*_*b*_ large with *b* constant, the expressions for the first two moments of RNAP at every gene segment and of mature RNA from Eqs. (2) and (4), respectively, simplify to

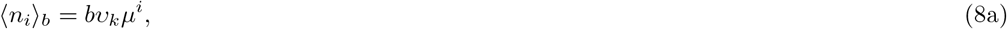

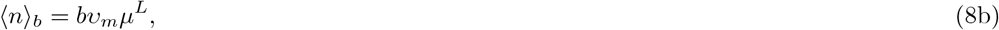

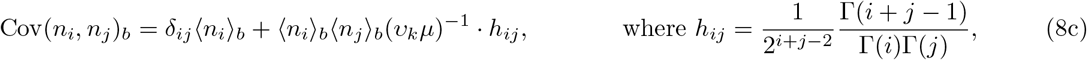

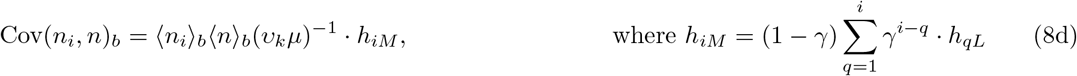

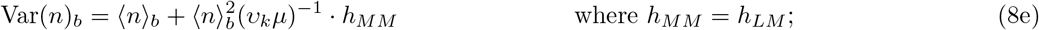

here, the subscript *b* denotes the moments in the bursty limit. Moreover, *υ*_*k*_ = *s*_*u*_/*k*, *υ*_*m*_ = *s*_*u*_/*d*_*m*_, and *h*_*ij*_ = *f*_*ij*_ | _*α*→0_ denotes the simplified function *f*_*ij*_, as 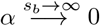 in that limit.

To test the accuracy of our theory, in Fig. 2 we compare our analytical expressions for the mean of local RNAP numbers, as well as for various measures of local RNAP fluctuations – the coefficient of variation CV, the Fano factor FF, and the Pearson correlation coefficient CC – with those calculated from stochastic simulation using Gillespie’s algorithm (SSA) [23]. Since we use parameters measured for a gene that demonstrates bursty expression (PDR5) [4], we test the accuracy of both the exact theory from Eqs. (2) and (4) and the approximate expressions given in Eq. (8). The perfect agreement between exact theory (solid lines) and simulation (dots) verifies the correctness of our analysis. The approximate theory (dashed lines) provides a reasonably good approximation; the mismatch can be decreased if the degree of burstiness is increased, i.e. by increasing the mean burst size *b* = *r*/*s*_*b*_. The following interesting observations can be made from these figures: (i) if the rate of premature detachment is greater than zero, then the mean of local RNAP decreases monotonically with the distance *i* from the promoter according to a power law, whereas that mean is constant along the gene if there is no premature detachment, as expected; (ii) the size of RNAP fluctuations, as measured by CV, decreases with *i* for small premature detachment rates, but increases with *i* for sufficiently large values of the detachment rate; (iii) the Fano factor approaches 1 – the value of FF for a Poissonian distribution – as *i* increases, which is due to the dispersal of the burst as stochastic elongation proceeds; (iv) the correlation coefficient between the local RNAP on two neighboring gene segments decreases monotonically with *i*, which is exacerbated by premature detachment and is a direct result of the stochasticity inherent in the elongation process.

**Figure 2:**
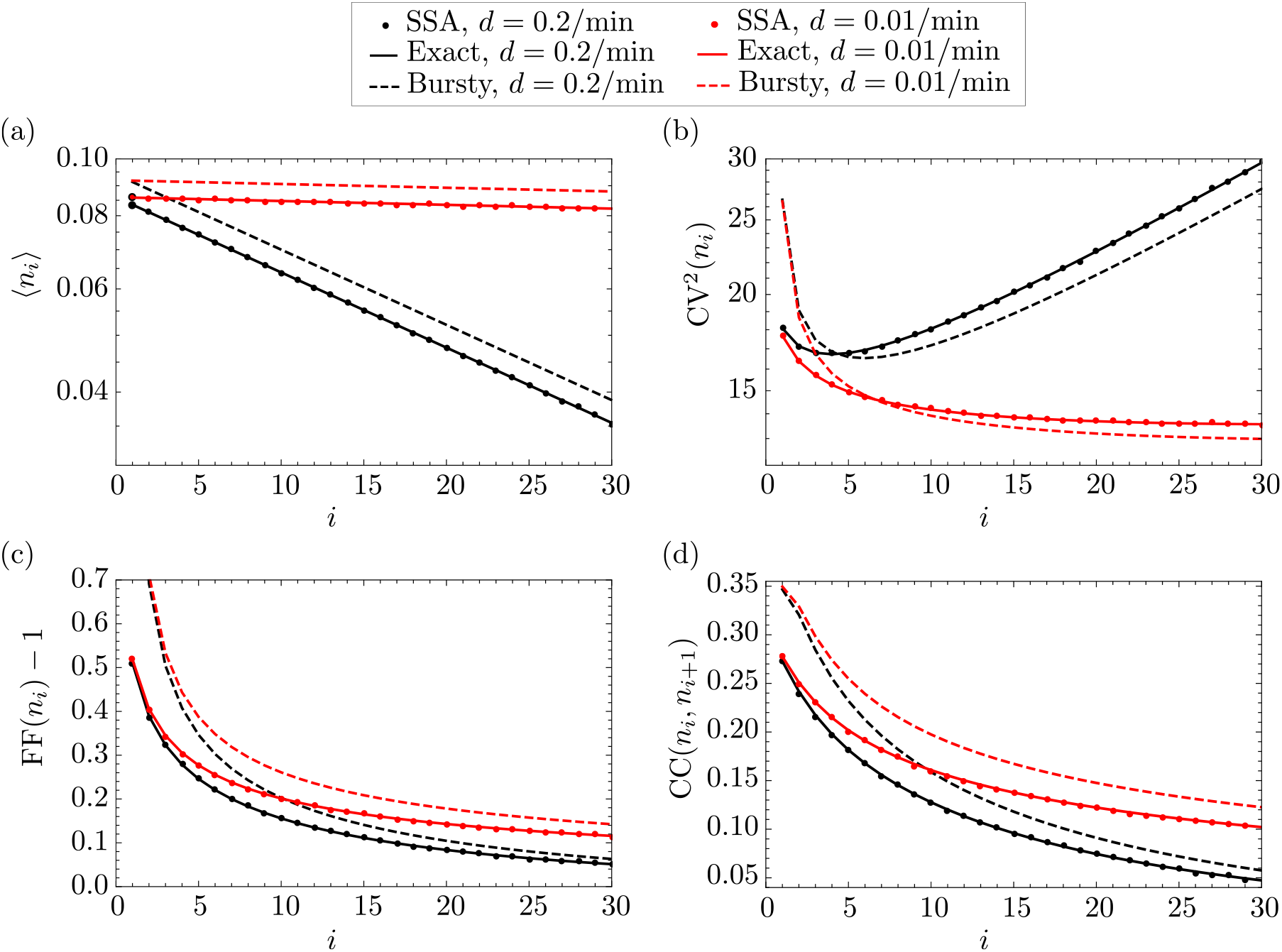
First and second moments of the distribution of local RNAP for the gene PDR5 in yeast, which demonstrates bursty expression. In panels (a), (b), (c), and (d), we show the dependence of the mean, coefficient of variation squared, Fano factor, and Pearson correlation coefficient, respectively, of local RNAP fluctuations on gene segment *i*, as predicted by our exact theory (Eqs. (2) and (4); solid lines), the approximate theory in the bursty limit (Eq. (8); dashed lines), and simulation via Gillespie’s stochastic simulation algorithm (SSA; dots), respectively. The parameters are fixed to *s*_*u*_ = 0.44/min, *s*_*b*_ = 4.7/min, and *r* = 6.7/min, which are characteristic of the PDR5 gene in yeast, as reported in Supplemental Table 2 of [4]. The number of gene segments is arbitrarily chosen to be *L* = 30. The total elongation time 〈*T*〉 = 4.5 min is also reported for PDR5, described as the synthesis time and denoted by *τ* in [4]. The elongation rate by definition takes the value of the ratio *k* = *L*/〈*T*〉 − *d* ≈ *L*/〈*T*〉, since *d* ≪ *k*. The detachment rate *d* is arbitrarily chosen to be *d* = 0.01/min (red lines and dots) or *d* = 0.2/min (black lines and dots). Note that, for the SSA, moments are calculated from one long trajectory with a few million time points, sampled at unit intervals.

The observation in (iii) can be explained in detail as follows. When the detachment rate is zero, a burst of RNAPs rapidly bind to the promoter, leading to large fluctuations near that site; however, thereafter each RNAP moves distinctly from all others due to stochastic elongation. Hence, the burst is gradually dispersed as elongation proceeds, which implies a decrease in the variance of fluctuations with increasing *i*. When the detachment rate is non-zero, then the same effect is at play; however, the increase in the variance of fluctuations along the gene is now counteracted by the decrease of mean RNAP numbers, which leads to two types of behaviour: for small *i*, CV decreases with *i*, since the variance dominates over the mean, while for large *i*, the opposite occurs and CV increases with *i*.

##### Constitutive limit

The other common parameter regime is that of constitutive gene expression, where the gene spends most of its time in the active state and transcription is continuous, which corresponds to the limit of very small *s*_*b*_. In that limit, the expressions from Eqs. (2) and (4) simplify to

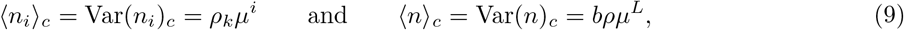

while the covariances Cov(*n*_*i*_, *n*_*j*_)_*c*_ and Cov(*n*_*i*_, *n*)_*c*_ between the species are zero; here, the subscript *c* denotes the constitutive limit. This drastic simplification reflects the fact that, in the constitutive limit, the distributions of mature RNA and local RNAP are Poissonian: as the regulatory network is effectively given by ∅ → *P*_1_ → *P*_2_ → ∆ *P*_*L*_ → *M* → ∅ then, the result follows directly from the exact solution provided in [24].

To further test the accuracy of our theory, in Fig. (3) we compare our analytical expressions for the mean of local RNAP numbers, as well as for various measures of local RNAP fluctuations, with those calculated from stochastic simulation using Gillespie’s algorithm, where we use parameters measured for a gene that demonstrates constitutive expression (DOA1) [4]. As before, we test the accuracy of both the exact theory given by Eqs. (2) and (4) and the approximate expressions from Eq. (9). Unsurprisingly, we observe agreement between exact theory (solid lines) and simulation (dots); the mismatch between our approximate theory and simulation is due to the fact that the gene does not spend 100% of its time in the active state – the true constitutive limit – but, rather, *s*_*u*_/(*s*_*u*_ + *s*_*b*_) ≈ 85%. The local mean RNAP number decreases with distance from the promoter, as was the case for bursty expression in the previous subsubsection, which is to be expected. The various measures which depend on the second moments are, however, considerably different: CV increases monotonically with *i*, independently of the rate of premature detachment, while FF and CC are very close to 1 and zero, respectively; moreover, the latter two measures practically show very little variation along the gene. The lack of transcriptional bursting explains all these effects in a straightforward fashion.

**Figure 3:**
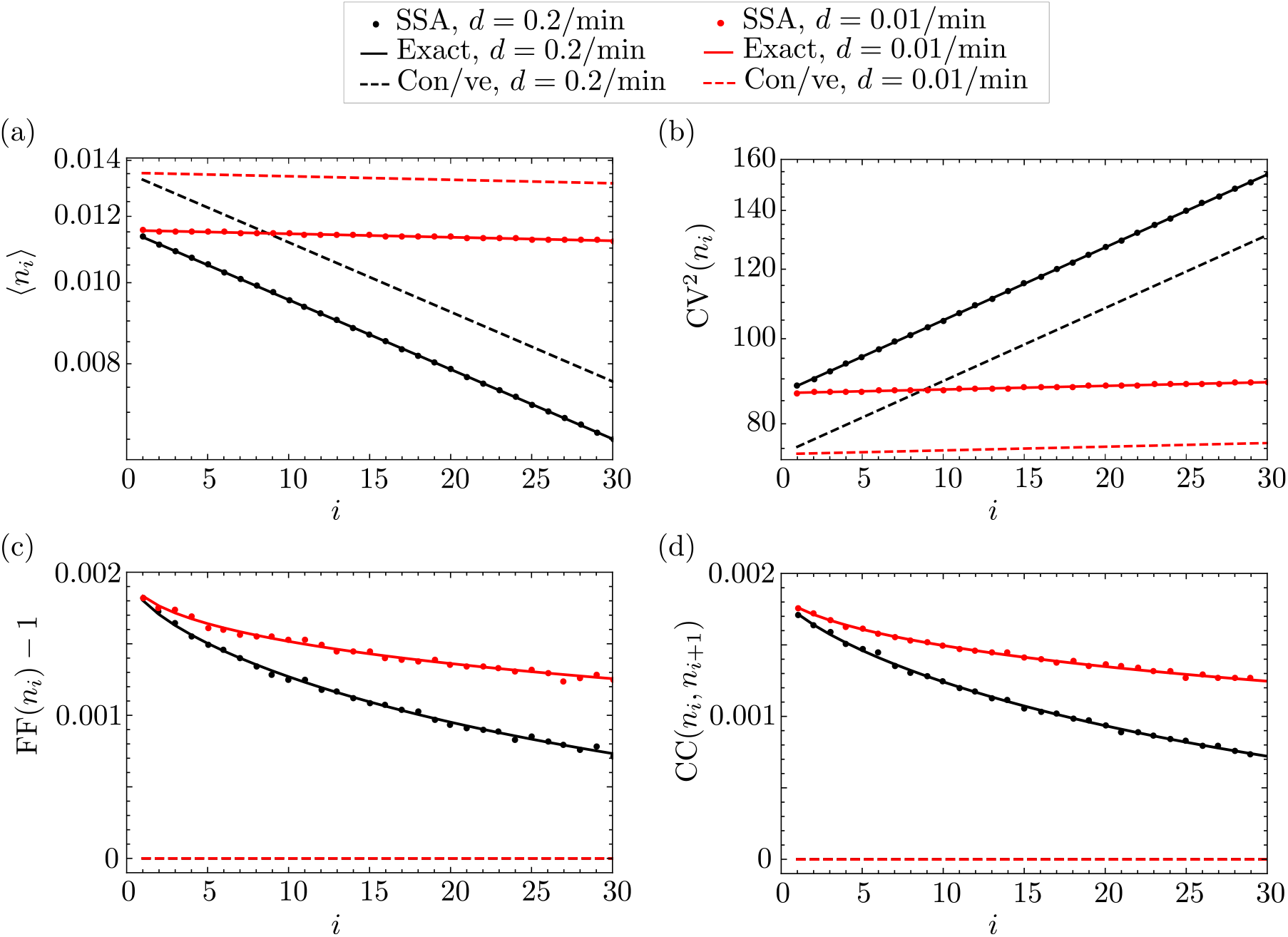
First and second moments of the distribution of local RNAP for the gene DOA1 in yeast, which demonstrates constitutive expression. In panels (a), (b), (c), and (d), we show the dependence of the mean, coefficient of variation squared, Fano factor, and Pearson correlation coefficient, respectively, of local RNAP fluctuations on gene segment *i*, as predicted by our exact theory (Eqs. (2) and (4); solid lines), the approximate theory in the bursty limit (Eq. (8); dashed lines), and simulation via Gillespie’s stochastic simulation algorithm (SSA; dots), respectively. The parameters are fixed to *s*_*u*_ = 0.7/min, *s*_*b*_ = 0.12/min and *r* = 0.14/min, which are characteristic of the DOA1 gene in yeast, as reported in Supplemental Table 2 of [4]. The number of gene segments is arbitrarily chosen to be *L* = 30. The total elongation time 〈*T*〉 = 2.9 min is also reported for DOA1, described as the synthesis time and denoted by *τ* in [4]. The elongation rate by definition takes the value of the ratio *k* = *L*/〈*T*〉 − *d* ≈ *L*/〈*T*〉, since *d* ≪ *k*. The detachment rate *d* is arbitrarily chosen to be *d* = 0.01/min (red lines and dots) or *d* = 0.2/min (black lines and dots). Note that, for the SSA, moments are calculated from one long trajectory with a few billion time points, sampled at unit intervals.

Finally, we remark that the accuracy of our expressions for the mean and variance of mature RNA, as given in Eq. (2) and (4), are verified by SSA simulation in Figs. 4(a) and (b) for parameters typical of the bursty PDR5 gene. The meaning of the dependence of descriptive statistics on *L* is discussed in the next section.

**Figure 4:**
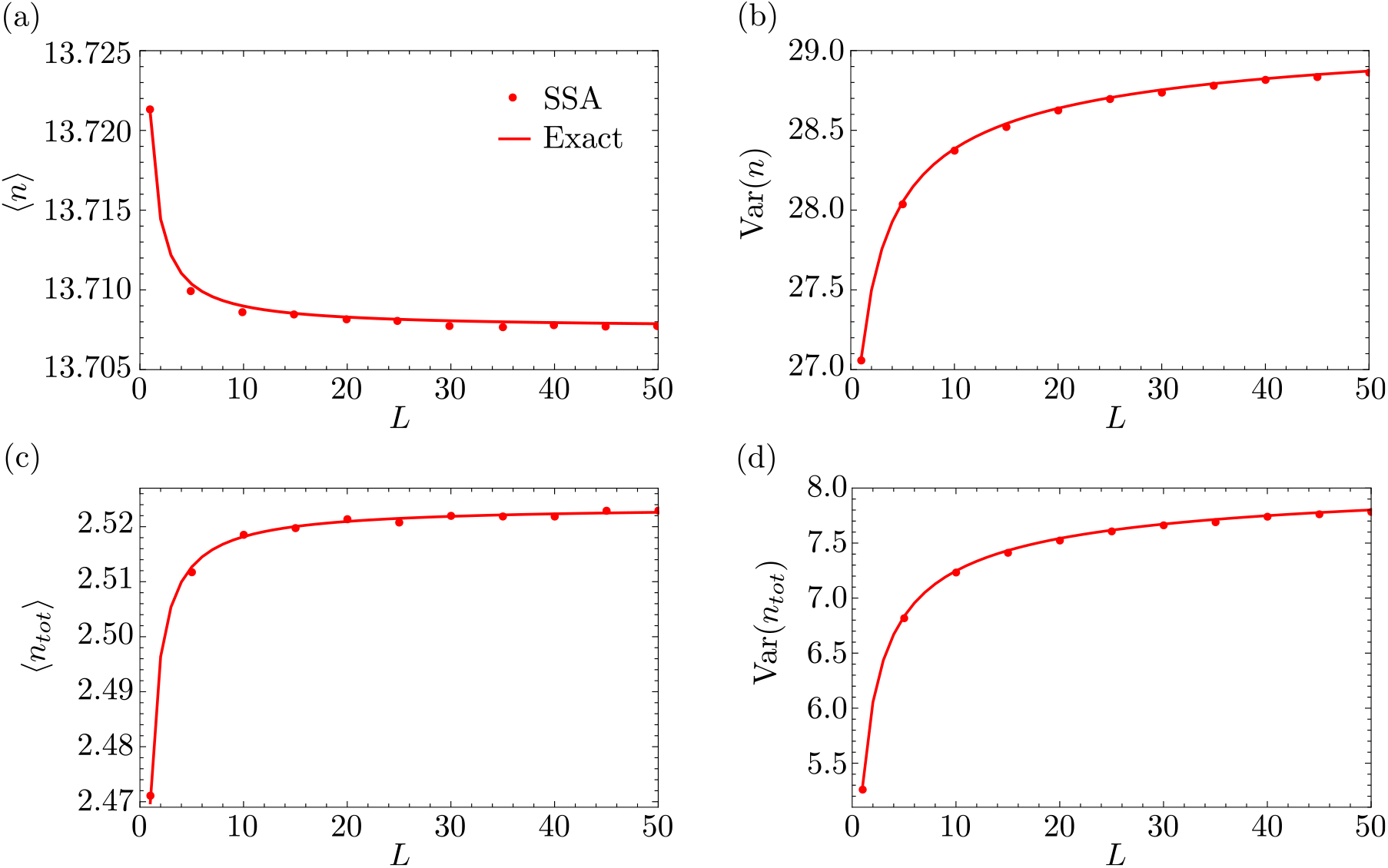
Mean and variance of the distributions of mature RNA and total RNAP for the PDR5 gene in yeast. In panels (a) and (b), we show the dependence of the moments of mature RNA fluctuations on the number of gene segments *L*, as predicted by our theory (Eqs. (2) and (4); solid lines) and SSA (dots). In panels (c) and (d), we show the dependence of the moments of total RNAP on *L*, as predicted by our exact theory (Eq. (10); solid lines) and SSA (dots). The parameters *s*_*u*_, *s*_*b*_, *r*, and 〈*T*〉 are characteristic of the PDR5 gene, and are the same as in Fig. 2. The premature detachment rate is chosen to be *d* = 0.01/min; the elongation rate is then given by *k* ≈ *L*/〈*T*〉. The degradation rate of mature RNA is *d*_*m*_ = 0.04/min, which is chosen such that the mean mature RNA is roughly consistent with that reported in Fig. 6(b) of [4]. Note that, for the SSA, moments are calculated from one long trajectory with a few billion time points, sampled at unit intervals.

### 2.3 Closed-form expressions for moments of total RNAP

While local RNAP fluctuations are measurable in experiment, as discussed in the Introduction, measurements of total RNAP on a gene are typically reported. Hence, in this section, we briefly discuss descriptive statistics of total RNAP fluctuations.

Recalling that *n*_*i*_ is the number of RNAP molecules on the *i*-th gene segment, the total number of RNAPs on the gene – arbitrarily divided into *L* segments – is given by 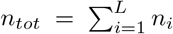. Given Eq. (2) and (4), the steady-state mean 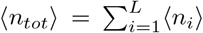 and the steady-state variance 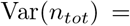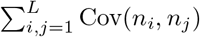 of the total RNAP distribution are given by

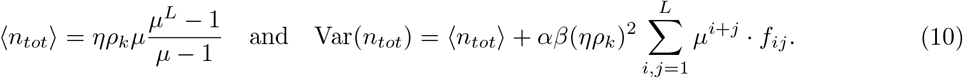

For a detailed derivation of the variance in Eq. (10), we refer to Appendix C. These expressions for the mean and variance of the total RNAP distribution simplify in the bursty and constitutive limits, as can be seen in Appendix D. The accuracy of Eq. (10) is tested by comparing against stochastic simulation with SSA in Figs. 4(c) and (d). Both mean and variance are seen to increase monotonically with the number of gene segments *L*, as we keep the mean elongation time constant; the mean shows very little dependence on *L*, while the dependence of the variance is more pronounced. We recall that, while the parameter *L* is arbitrary in principle, it actually determines the size of fluctuations in the elongation time. Since that time is the sum of *L* independent exponential variables with mean 1/(*k* + *d*) each, it follows that the distribution of the elongation time *T* is Erlang with mean 〈*T*〉 = *L*/(*k* + *d*) and coefficient of variation squared equal to 1/*L*. Hence, the larger *L* is, the narrower is the distribution of *T* and the more deterministic is elongation itself. Thus, Figs. 4(c) and (d) predict that the mean and variance of total RNAP increase rapidly with decreasing fluctuations in the elongation time *T*. It hence follows that models in which the elongation rate is assumed to be exponentially distributed [10], which correspond to the case where *L* = 1 in our model, underestimate the size of nascent RNA fluctuations.

### 2.4 Special case of deterministic elongation

Next, we derive expressions for the descriptive statistics of total RNAP and mature RNA in the limit of large *L* taken at constant mean elongation time, which corresponds to deterministic elongation. As is shown in Fig. 4, these statistics converge quickly to the ones obtained in the large-*L* limit; hence, the resulting limiting expressions are likely to be useful across a variety of genes.

#### Moments of total RNAP distribution

We define the non-dimensional parameters δ_*g*_ = *τ*_*g*_/*τ*_*d*_, *T*_*g*_ = 〈*T*〉/*τ*_*g*_, and *T*_*d*_ = 〈*T*〉/*τ*_*d*_, which correspond to the ratio of the gene timescale and the polymerase detachment timescale, the ratio of the mean elongation timescale and the gene timescale, and the ratio of elongation timescale and the polymerase detachment timescale, respectively; here, *τ*_*d*_ = 1/*d*, as before. Substituting *k* ↦ *L*/〈*T*〉 − *d* into Eq. (10) and taking the limit of deterministic elongation, i.e. letting *L* → ∞ at constant 〈*T*〉, we obtain the following expressions for the mean, variance, and CV^2^ of total RNAP:

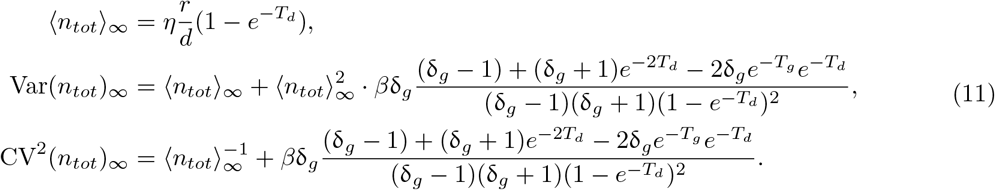

Here, the subscript ∞ denotes the limit of *L* → ∞. A detailed derivation of the variance in Eq. (11) can be found in Lemma C.1 of Appendix C.

In the special case when RNAP does not prematurely detach from the gene, i.e. for *d* = 0, the expressions in Eq. (11) simplify to

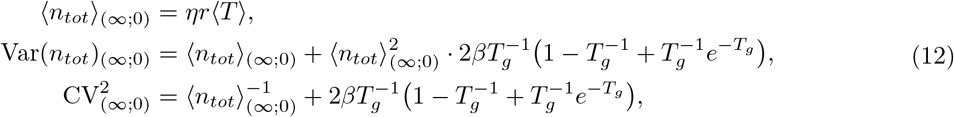

where the subscript (∞; 0) denotes the limit of (*L, d*) → (∞, 0). The expressions in Eq. (12) have been previously reported in [7], where they were derived using queuing theory. Hence, our expressions in Eq. (11) constitute a generalization of known results, by further taking into account premature detachment of RNAP from the gene.

Eq. (12) shows that the coefficient of variation squared of total RNAP, denoted by 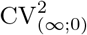, can be written as the sum of two terms: (i) the inverse of the mean which is expected if the distribution of total RNAP is Poissonian, and (ii) a term that increases with increasing *β* and decreasing *T*_*g*_. Hence, the latter term provides a measure for the deviation of the total RNAP distribution from a Poissonian. In particular, it shows that the deviation is significant in genes for which (i) the fraction of time spent in the inactive state is large (large *β*), and (ii) the elongation time is much shorter than the switching time between the active and inactive states (small *T*_*g*_).

#### Moments of mature RNA distribution

Similarly, in the limit of deterministic elongation, it is straightforward to show that the expressions for the mean and variance of the distribution of mature RNA given by Eqs. (2) and (4) reduce to

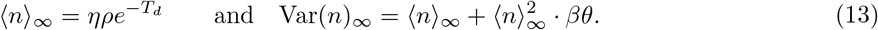

These expressions can be further simplified in the special cases of no premature detachment to read

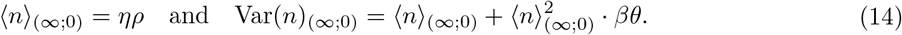

Note that the mean and variance are precisely the same as would be obtained from the telegraph model, for which the corresponding Fano factor in the bursty limit is given by Eq. (16) below. Hence, we anticipate that, in the limit of no premature detachment and deterministic elongation, the distribution of mature RNA from our transcription model is the same as the distribution obtained from the coarser telegraph model. A formal proof of that claim will be given in Section 3.

#### Relationship between Fano factors of total RNAP and mature RNA

Specifying to the case of no premature detachment, it is interesting to note that in the bursty limit, i.e. for *r, s*_*b*_ → ∞ at constant mean burst size *b* = *r/s*_*b*_ in Eq. (12), the Fano factor of total RNAP is given by

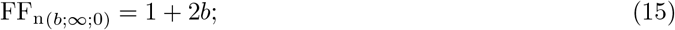

see also Eq. (D.3) in Appendix D. Here, the subscript n denotes nascent RNA (total RNAP). Eq. (15) is in contrast to the Fano factor of mature RNA in the same bursty limit:

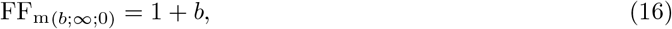

see Eq. (D.8) in Appendix D, where the subscript *m* denotes mature RNA. (Note that FF_m(*b*;∞;0)_ also equals the Fano factor of the telegraph model in the same bursty limit [25].) Hence by comparing Eqs. (15) and (16) we can deduce the following for bursty expression: (i) if one uses the telegraph model to estimate the mean transcriptional burst size from total RNAP data where the elongation time is deterministic, then the mean burst size will be overestimated by a factor of two – in other words, the implicit assumption that the elongation time is exponentially distributed is inadequate; (ii) fluctuations in total RNAP (nascent RNA) deviate more from Poisson statistics, for which the Fano factor equals one, than fluctuations in mature RNA.

More generally, if we do not enforce the bursty limit, then we find the following relationship between the Fano factors of total RNAP and mature RNA, which are calculated from Eqs. (12) and (14), respectively:

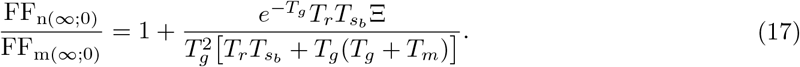

Here,

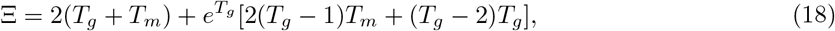

while *T*_*g*_ = (*s*_*u*_ + *s*_*b*_) 〈*T*〉, *T*_*r*_ = *r*〈*T*〉, *T*_*m*_ = *d*_m_ 〈*T*〉, and 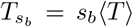 are non-dimensional parameters representing the ratio of the mean elongation time to the timescales of promoter switching, initiation, decay of mature RNA, and gene deactivation, respectively. From Eq. (17), we deduce that FF_n(∞;0)_ > FF_m(∞;0)_ if and only if Ξ > 0. From the contour plot of Ξ in Fig. 5, one can deduce that

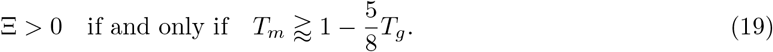

**Figure 5:**
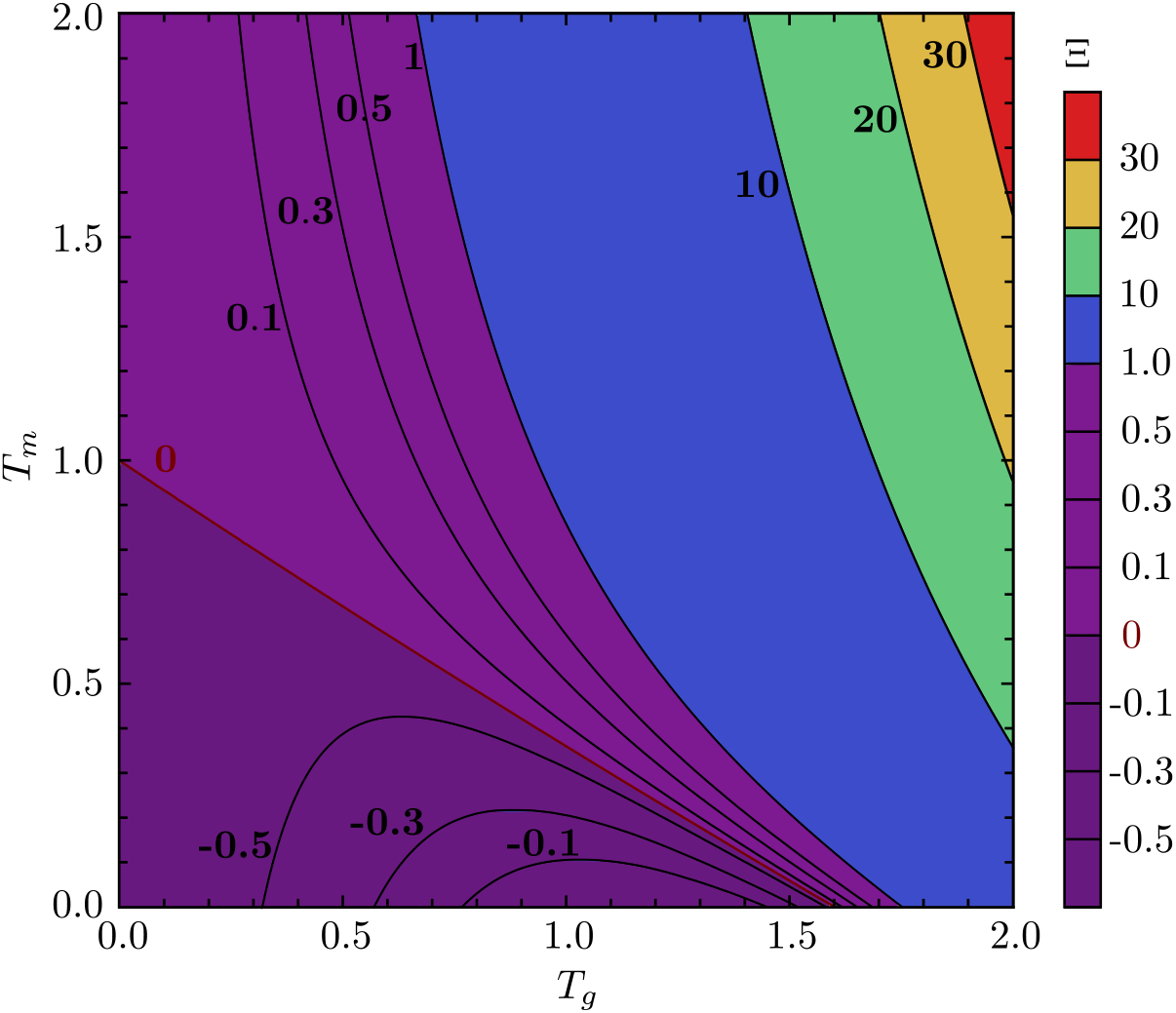
Comparison between the Fano factors of nascent and mature RNA. Contour plot showing the variation of Ξ – a measure of the difference between the two Fano factors which is defined in Eq. (18) – with the non-dimensional parameters *T*_*g*_ and *T*_*m*_ which denote the ratio of the mean elongation timescale to that of the timescales of promoter switching and of mature RNA decay, respectively. As can be appreciated from Eq. (17), Ξ is positive if the Fano factor of nascent RNA is larger than that of mature RNA and negative if the reverse is true. The line 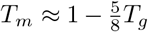, where Ξ = 0, shows where the two Fano factors are identical.

Hence, the Fano factor of nascent RNA is larger than that of mature RNA if and only if the above (approximate) condition is satisfied. In the bursty limit, *T*_*g*_ → ∞ due to *s*_*b*_ → ∞ which, together with *T*_*m*_ > 0, implies that Eq. (19) holds; the condition is also satisfied if promoter switching is very fast compared to elongation. In contrast, if *T*_*m*_ < 1 and *T*_*g*_ < 1, then it is possible to have the opposite scenario where the Fano factor of mature RNA is larger than that of nascent RNA, which occurs for example if promoter switching and mature RNA decay are very slow compared to elongation.

#### Sensitivity of coefficient of variation of total RNAP and mature RNA

Since we have found explicit expressions for the first two moments of the distributions of total RNAP and of mature RNA, we can now estimate the sensitivity of the noise in each of those to small perturbations in the transcriptional parameters. Specifically, we calculate the logarithmic sensitivity (LS), which is also known as the relativity sensitivity, of the coefficient of variation to a parameter *s*, which is defined as Λ_*s*_ = (*s*/CV)(*∂*CV/*∂s*). (That definition implies that a 1% change in the value of the parameter *s* results in a change of Λ_*s*_% in CV.)

In Table 1b, we report the logarithmic sensitivity of the coefficient of variation (CV) of total RNAP fluctuations, which is obtained from Eq. (12), to perturbations in the parameters *s*_*u*_, *s*_*b*_, *r*, and 〈*T*〉. Similarly, in Table 1c, we report the logarithmic sensitivity of the coefficient of variation of mature RNA fluctuations from Eq. (14) to perturbations in the parameters *s*_*u*_, *s*_*b*_, *r*, and *d*_*m*_. In both cases, these sensitivities are calculated for parameter values estimated for five genes in yeast, as reported in [4]; see Table 1a.

**Table 1:**
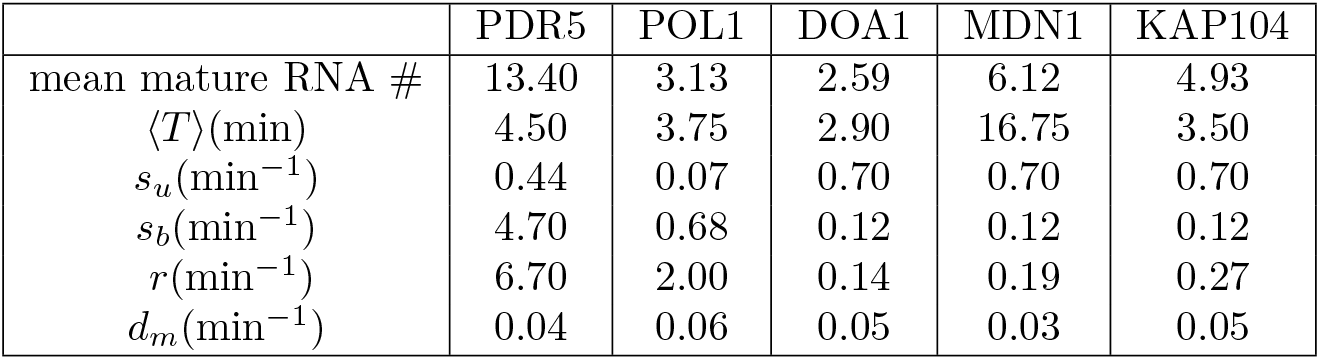
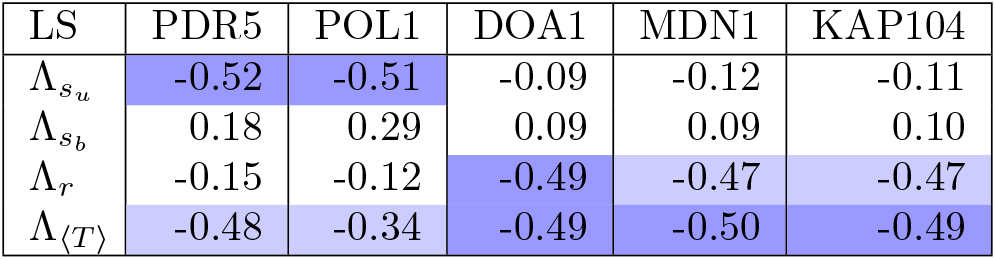
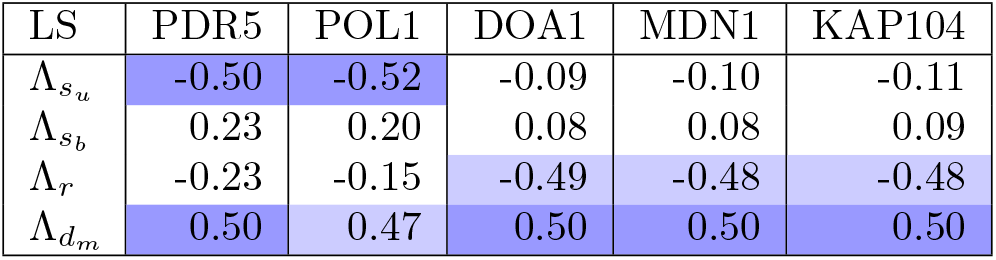
Logarithmic sensitivity (LS) of the coefficient of variation CV of total RNAP and mature RNA fluctuations for five genes in yeast; see Section 2.4 for a discussion. (a) Parameter values from Supplemental Tables 2 and 4 in [4]. The degradation rate *d*_*m*_ of mature mRNA is estimated from the reported mean number of mature RNA, the parameters *s*_*u*_, *s*_*b*_, *r*, and Eq. (14) for the mean. (b) Logarithmic sensitivity of CV of total RNAP fluctuations. The most sensitive parameter and the next most sensitive one are marked in dark and light blue, respectively. (c) Logarithmic sensitivity of CV of mature mRNA fluctuations. The most sensitive parameter and the next most sensitive one are marked by dark and light blue, respectively.

The following observations can be made regarding the sensitivity of the noise in total RNAP fluctuations: (i) for the two genes PDR5 and POL1 which spend most of their time in the inactive state due to *s*_*b*_ ≫ *s*_*u*_, CV is most sensitive to changes in the parameters *s*_*u*_ and 〈*T*〉; (ii) for the genes DOA1, MDN1, and KAP104, which spend most of their time in the active state due to *s*_*u*_ ≫ *s*_*b*_, CV is most sensitive to changes in the parameters *r* and 〈*T*〉; (iii) the size of mature RNA fluctuations is found to be most sensitive to perturbations in *s*_*u*_ and *d*_*m*_ for PDR5 and POL1, and to perturbations in *r* and *d*_*m*_ for the other three genes. We furthermore note that for both total RNAP and mature RNA, *r* is the least sensitive parameter for the genes which are mostly inactive, whereas it is among the most sensitive parameters for genes that are mostly active.

## 3 Approximate distributions of total RNAP and mature RNA

Thus far, we have derived expressions for the first two moments of the distributions of total RNAP and mature RNA. Naturally, it would also be useful to derive closed-form expressions for the distributions themselves; such a derivation is, however, analytically intractable in general [24] due to the presence of the catalytic reaction *G*_*on*_ → *G*_*on*_ + *P*_1_, which models initiation of the transcription process. Still, there are two special cases where analytical distributions are known: (i) when the elongation time is considered to be fixed, which corresponds to our model with *L* → ∞ at constant 〈*T*〉 [9]; (ii) when the elongation time is exponentially distributed, corresponding to our model with *L* = 1, in which case the distribution of total RNAP is identical to the one which is derived from the telegraph model [6, 25]. While one may argue that the analytical distribution of RNAP for deterministic elongation times may well approximate the stochastic (finite-*L*) case, the issue remains that the exact solution is not given in terms of simple functions unless promoter switching is slow compared to initiation, elongation and termination, in which case the solution reduces to a weighted sum of two Poisson distributions [9]. Hence, it is generally very difficult to apply in practice, such as to infer parameters from data using a Bayesian approach. Moreover, to our knowledge, no exact solutions are known for the distribution of mature RNA in our model. In this section, we aim to devise a simple approximation for the distribution of total RNAP numbers in terms of the Negative Binomial (NB) distribution; these simple distributions have shown great flexibility in describing complex gene expression models with a large number of parameters [10]. Finally, we will obtain the distribution of mature RNA in the limit of deterministic elongation by means of singular perturbation theory.

### 3.1 Approximation of total RNAP distribution

We approximate the distribution of total RNAP transcribing the gene via a Negative Binomial distribution, as follows.

The mean and variance of the Negative Binomial distribution NB(*q, p*) are given by *pq*/(1 − *p*) and *pq/*(1 − *p*)^2^, respectively. By assuming that these are equal to the exact mean and variance, respectively, of the total RNAP distribution, see Eq. (10), we obtain effective values for the parameters *p* and *q*:

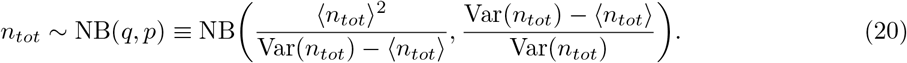

In Fig. 6, we show a comparison between the distributions of total RNAP obtained from the SSA (dots) and the Negative Binomial approximation in Eq. (20 (lines). Our results are presented for two different values of the number of gene segments: *L* = 1 (exponentially distributed elongation time; left column) and *L* = 50 (quasi-deterministic elongation time; right column). Additionally, we rescale our gene inactivation rate as *s*_*b*_ ↦ *s*_*b*_*ϵ*, and we present results for three different values of the parameter *ϵ*: 10^−3^, the constitutive limit of the gene being mostly in the active state (top row); 10^−1^, with the gene spending almost equal amounts of time in the active and inactive states, with *s*_*b*_ ≈ *s*_*u*_ (middle row); and 1, the bursty limit, where the gene spends most of its time in the inactive state (bottom row).

**Figure 6:**
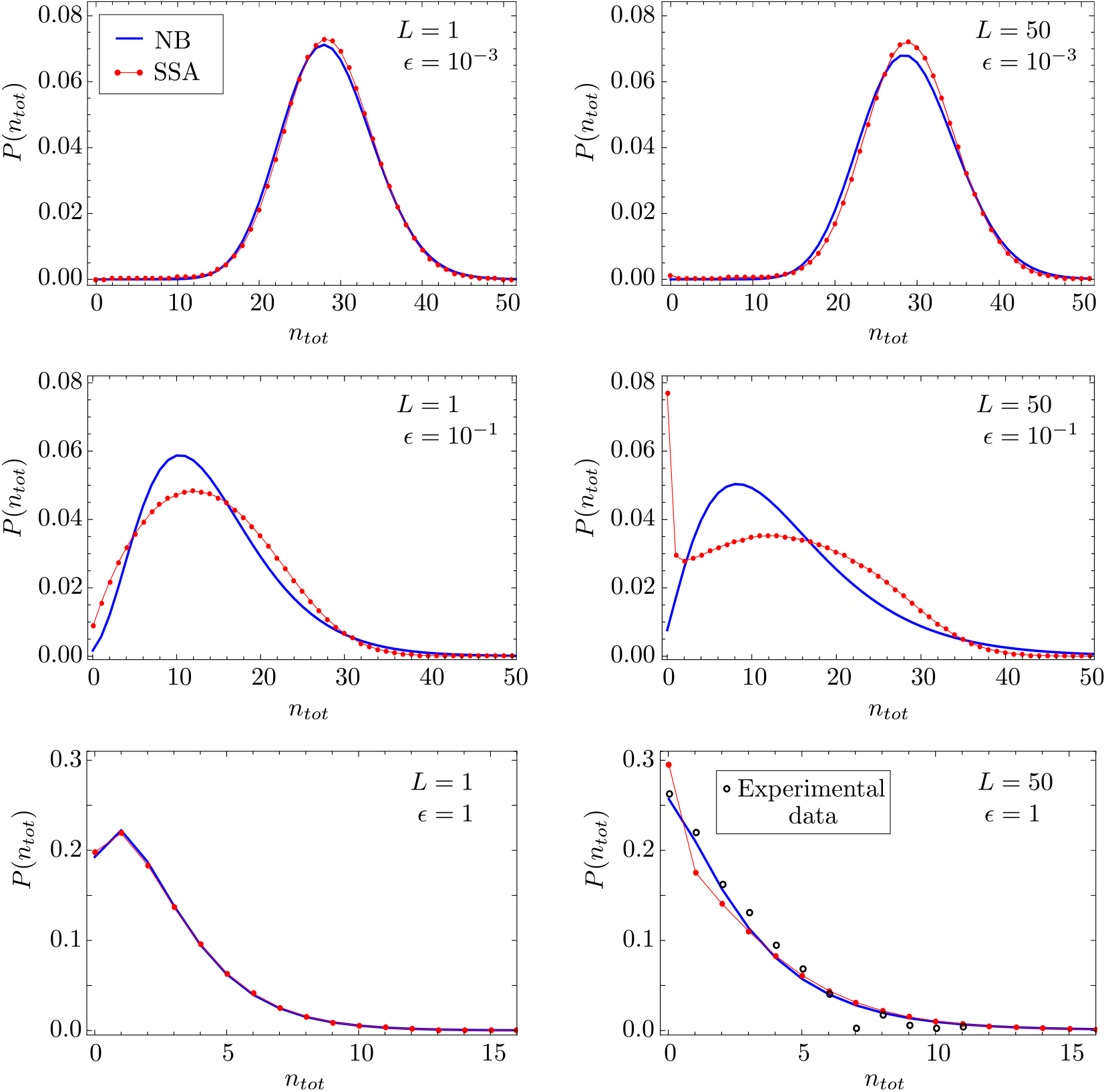
Steady-state distribution of total RNAP and its approximation by a Negative Binomial distribution. We compare the approximation from Eq. (20) (blue lines) with the distribution of total RNAP obtained from stochastic simulation (SSA; red dots). With the exception of *s*_*b*_, the parameters are for the PDR5 gene in yeast, and are hence the same as in Fig. 2, with *d* = 0.01/min. Results are presented for two different values of *L*, corresponding to an exponentially distributed elongation time (*L* = 1) and a quasi-deterministic elongation time (*L* = 50); *k* is rescaled such that the two have the same mean elongation time. Additionally, we rescale the gene inactivation rate via *s*_*b*_ ↦ *s*_*b*_*ϵ*, where *ϵ* = 10^−3^, 10^−1^, 1, corresponding to constitutive, general, and bursty expression, respectively. (Here, general expression is neither clearly constitutive nor bursty, since the gene spends roughly equal amounts of time in the inactive and active states.) Note that *ϵ* = 1 results in a distribution of nascent RNA that is consistent with that measured for PDR5; the experimental data from Fig. 6(b) of [4] is plotted for comparison. The Negative Binomial approximation is found to be accurate in the limits of constitutive and bursty expression (top and bottom rows), independently of *L*.

We can make several observations, as follows. For both *L* = 1 and *L* = 50, the Negative Binomial approximation performs well for bursting and constitutive expression (top and bottom rows), whereas it is appreciably poor when expression is in between those two limits (middle row). Intuitively, that observation can be explained via the following reasoning. In the limits of the gene being mostly in the active state (constitutive expression) or the inactive state (bursty expression), the distribution of total RNAP is necessarily unimodal. However, when the gene spends a considerable amount of time in each state, the distribution is the sum of two conditional distributions which can manifest either as bimodality or as a wide unimodal distribution, neither of which can be captured by a Negative Binomial distribution. Assuming bursty expression, the Negative Binomial distribution is a more accurate approximation to the distribution obtained from SSA for *L* = 1 than it is for *L* = 50; the resaon is that *L* = 1 corresponds to the telegraph model [25], in which case it can be proven analytically that the distribution reduces to a Negative Binomial in the limit of bursty expression. For constitutive expression, the Negative Binomial approximation is equally good for *L* = 1 and *L* = 50, as the distribution is necessarily Poissonian then and as it is well known that a Negative Binomial distribution can approximate a Poissonian to a high degree of accuracy. In summary, our results hence indicate that Eq. (20) yields a good approximation for the total RNAP distribution of bursty and constitutively expressed genes.

We also note from Fig. 6 that the comparison between the SSA distributions for *L* = 1 and *L* = 50, with equal mean elongation times, highlights the importance of modelling elongation with the correct distribution of elongation times for genes that are non-constitutive, i.e. for *ϵ* = 10^−1^ or *ϵ* = 1. In particular, if the elongation time is quasi-deterministic (*L* = 50), there appears to be a significant increase in the probability of observing zero total RNAP transcribing the gene compared to models with an exponentially distributed elongation time (*L* = 1).

### 3.2 Approximation of mature RNA distribution

Next, we apply singular perturbation theory to formally derive the distribution of mature RNA when the elongation time is deterministic.

We start by defining 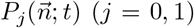 as the probability of the state 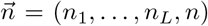 at time *t* while the gene is either active (0) or inactive (1). Note that *n*_*i*_ is the number of RNAPs on gene segment *i* for *i* = 1, …, *L*, while *n* is the number of mature RNAs. The time evolution of the probabilities 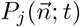 can be described by a system of coupled CMEs:

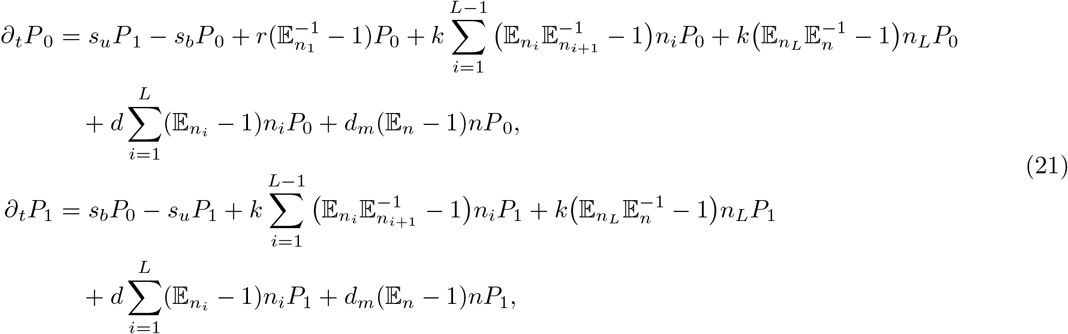

where 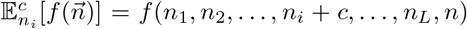, with *c* ∈ ℤ, denotes the standard step operator. We assume that the elongation rate *k* is faster than the degradation rate *d*_*m*_ of mature RNA, i.e. that *k*/*d*_*m*_ ≫ 1. Since *k* = *L*/ 〈*T*〉 − *d*, it follows that in the limit of deterministic elongation (*k* → ∞), i.e. for *L* → ∞ at constant mean elongation time 〈*T*〉, the condition *k*/*d*_*m*_ ≫ 1 is naturally satisfied.

In order to find an analytical expression for the propagator probabilities 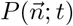 which satisfies the system of CMEs in Eq. (21), we define the probability-generating function as *F* = Σ_*j*_ *F*_*j*_, with 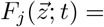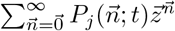; here, 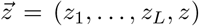 is a vector of variables corresponding to the state 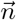. Given the equations for 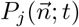 from Eq. (21), we obtain the following systems of PDEs for the corresponding generating functions 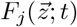:

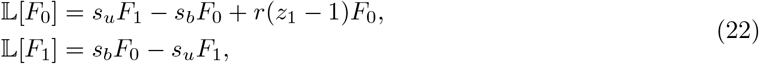

where

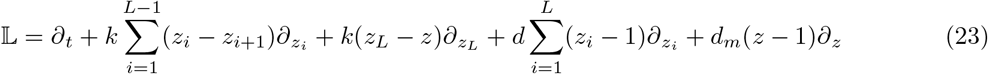

is a differential operator acting on the generating functions *F*_0_ and *F*_1_. Eq. (22) represents a system of coupled, linear, first-order PDEs. Now, we introduce the new variables *u*_*i*_ = *z*_*i*_ − 1 (*i* = 1, …, *L*) and *u* = *z* − 1 to rewrite Eq. (22) as

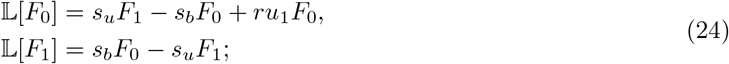

here, the operator in Eq. (23) now takes the form

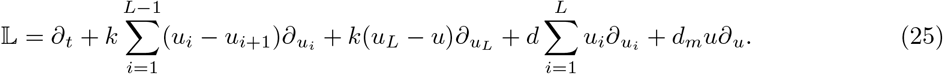

In order to find an analytical solution to Eq. (24), we rescale all rates and the time variable by the decay rate of mature RNA; then, we apply the method of characteristics, with *s* being the characteristic variable. The first characteristic equation gives *d*_*m*_(*dt/ds*) = 1, with solution *s* ≡ *t*′ = *d*_*m*_*t*; hence, we can use the variable *t′* as the independent variable and thus convert the system of PDEs in Eq. (24) into a characteristic system of ODEs,

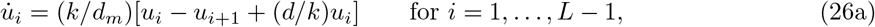

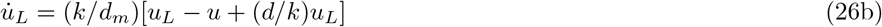

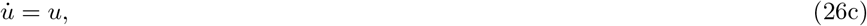

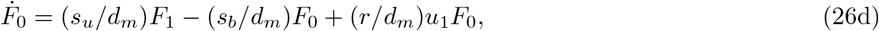

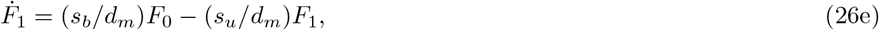

where the overdot denotes differentiation with respect to *t*′. The existence of an integral-form solution to Eq. (26) follows from the fact that the reaction scheme in Fig. 1 contains first-order reactions only. Under the assumption that *k* ≫ *d*_*m*_, we define *ɛ* = *d*_*m*_/*k*; then, we apply Geometric Singular Perturbation Theory (GSPT) [26, 27], with 0 < *ɛ* ≪ 1 as the (small) singular perturbation parameter. We hence separate the system in Eq. (26) into fast and slow dynamics, which will allow us to find an asymptotic approximation for *F*_0_ and *F*_1_ in steady-state. A brief introduction to GSPT can be found in Appendix E. Given the above definition of *ɛ*, Eqs. (26a) and (26b), the governing equations for *u*_*i*_ in the ‘slow system’, become

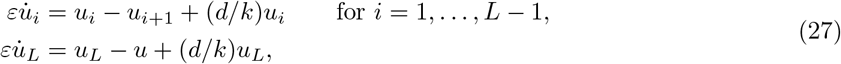

where *u*_*i*_ (*i*, …, *L*) are the fast variables and *u*, *F*_0_, and *F*_1_ are the slow ones. Setting *ɛ* = 0 in Eq. (27), we can express the variables *u*_*i*_ as *u*_*i*_ = *μ* · *u*_*i*+1_, with *μ* = *k*/(*k* + *d*) for *i* = 1, …, *L*. Finally, we write the variable *u*_1_ as *u*_1_ = *μ*^*L*^ · *u*. Next, given Eq. (26c), we apply the chain rule, with *dt*′ ≡ *du* · *u*, to rewrite Eqs. (26d) and (26e) as

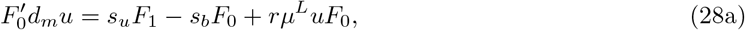

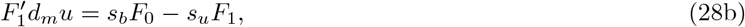

where the prime now denotes differentiation with respect to *u*. Solving Eq. (28a) for *F*_1_ and substituting the result into Eq. (28b), we obtain the second-order ODE

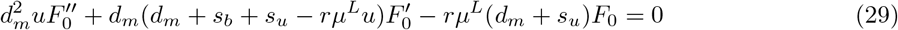

for *F*_0_(*u*). Eq. (29) is a confluent hypergeometric differential equation (Kummer’s equation) [28] which admits the solution

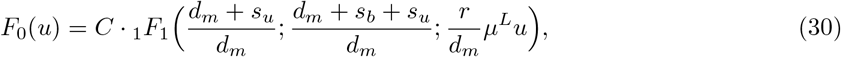

where _1_*F*_1_ denotes the confluent hypergeometric function; here, we consider only one of two independent fundamental solutions of Kummer’s differential equation, as we are seeking a solution in steady-state where the variable *u* is bounded. The constant *C* in Eq. (30) is a constant of integration that is determined from the normalization condition on the full generating function: *F* = *F*_0_ + *F*_1_. From Eq. (28), one finds that *F* satisfies

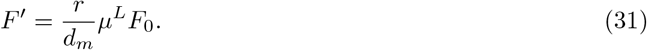

Making us of Eq. (31) and applying the normalization condition *F*|_*u*=0_ = 1, we find that the generating function at steady-state reads

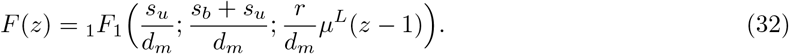

The probability distribution *P*(*n*) of mature RNA can be found from the formula

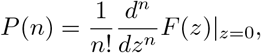

which yields the analytical expression

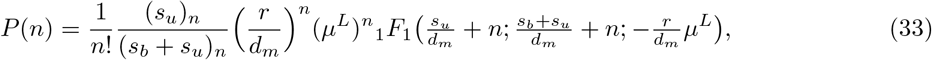

where (·)_*n*_ is the Pochhammer symbol, as before. Note that the mean and variance of mature mRNA, as calculated from the distribution in Eq. (33), agree exactly with Eqs. (2c) and (4f) in the limit of fast elongation rate (*k* → ∞). Note also that the solution in Eq. (33) depends on the parameter *μ*^*L*^, which represents the survival probability of an RNAP molecule, i.e. the probability that RNAP will not prematurely detach from the gene. Finally, we take the limit of deterministic elongation, letting *L* → ∞ at constant 〈*T*〉, which leads to

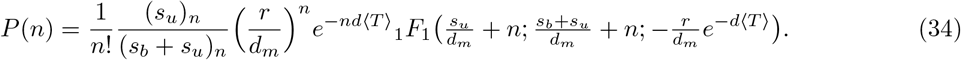

Note that in the limit of zero premature detachment (*d* = 0), Eq. (34) is precisely equal to the distribution of mature RNA predicted by the telegraph model, which is in wide use in the literature [25]. Hence, our perturbative approach can be seen as a means to formally derive the conventional telegraph model of gene expression starting from a more fundamental and microscopic model. In Fig. 7, we verify our analytical solution with stochastic simulation for two different genes in yeast. We also note that, for non-zero premature detachment (*d* ≠ 0), Eq. (34) is the steady-state solution predicted by the telegraph model, with parameter *r* renormalised to *re*^−*d*〈*T*〉^.

**Figure 7:**
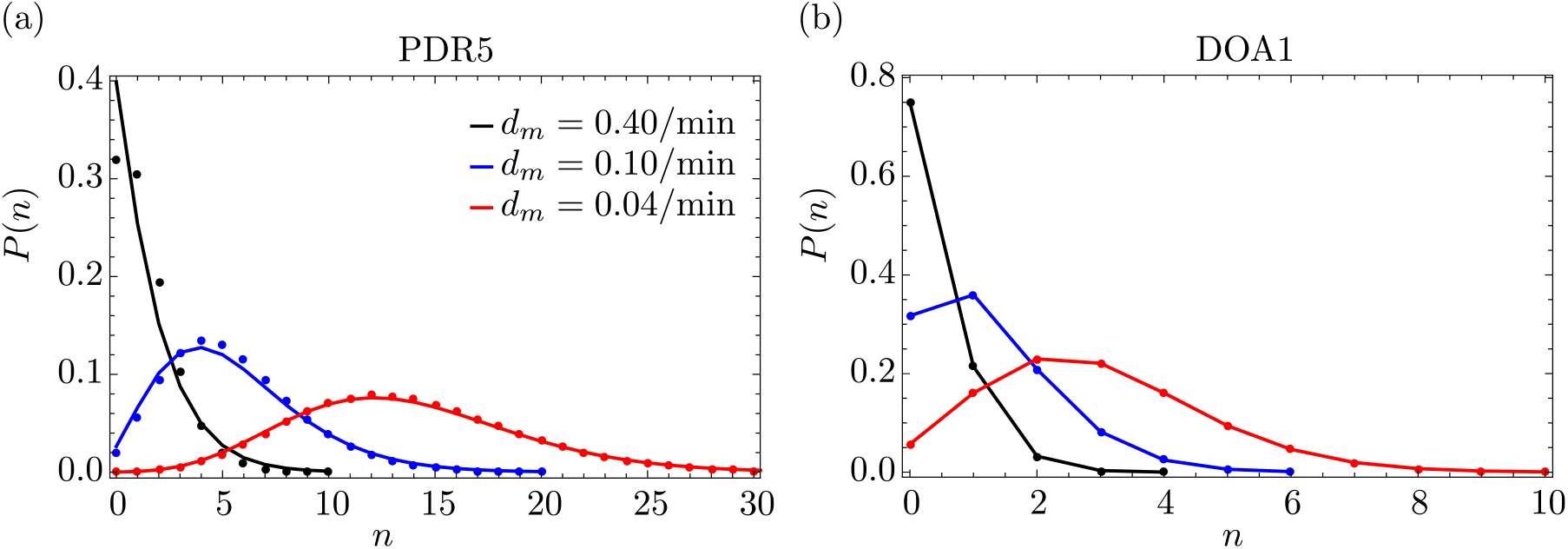
Steady-state distribution of mature RNA for two different genes in yeast. We compare the distribution obtained from the SSA (dots) to the perturbative approximation in Eq. (33) (solid lines) for two different genes. In panel (a), we consider the PDR5 gene, fixing the parameters as in Fig. 2: *s*_*u*_ = 0.44/min, *s*_*b*_ = 4.7/min, *r* = 6.7/min, *d* = 0.01/min, and 〈*T*〉 = 4.5 min. The degradation rate of mature RNA takes the values *d*_*m*_ = 0.04, 0.10, 0.40/min; note that the experimental value is *d*_*m*_ = 0.04/min. In panel (b), we consider the DOA1 gene, fixing the parameters as in Fig. 3: *s*_*u*_ = 0.7/min, *s*_*b*_ = 0.12/min, *r* = 0.14/min, *d* = 0.01/min, and 〈*T*〉 = 2.9 min. The degradation rate of mature RNA again takes the values *d*_*m*_ = 0.04, 0.10, 0.40/min; the experimental value is *d*_*m*_ = 0.05/min. For both genes, the agreement between SSA and our perturbative approximation increases with *k*/*d*_*m*_, as expected, since Eq. (33) is derived under the assumption that *k* ≫ *d*_*m*_. Note that the distribution is practically independent of *L*, since Eq. (33) depends on *L* only through *μ*^*L*^, which for small premature detachment rates *d* implies *μ*^*L*^ ≈ 1 for any *L*.

## 4 Statistics of fluorescent nascent RNA signal

Thus far, we have determined the statistics of the total number of RNAP transcribing the given gene; these are also the statistics of the number of nascent RNA molecules. However, in experiments using single-molecule fluorescence in situ hybridization (smFISH [9]), molecule numbers of nascent RNA cannot be directly determined. Rather, the experimentally measured RNA ‘abundance’ is the fluorescent signal emitted by oligonucleotide probes bound to the RNA. Since the length of the nascent RNA grows as RNAP moves away from the promoter, it follows that we must account for the increase in the fluorescent signal as elongation proceeds.

In this section, we take into account these experimental details to obtain closed-form expressions for the mean and variance of the fluorescent signal of local and total nascent RNA. We assume that the signal from nascent RNA on the *i*-th gene segment is given by *r*_*i*_ = (*v/L*)*in*_*i*_ for *i* = 1, …, *L*, where *v* is some experimental constant; the value of the parameter (*v/L*)*i* is increasing with *i*, which models the fact that the fluorescent signal becomes stronger as RNAP moves along the gene. The formula for the mean fluorescent signal at gene segment *i* is then given by 〈*r*_*i*_〉 = (*v/L*)*i*〈*n*_*i*_〉, where 〈*n*_*i*_〉 follows from Eq. (2b); the covariance of two fluorescent signals along the gene, *r*_*i*_ and *r*_*j*_ (*i, j* = 1, …, *L*), is given by Cov(*r*_*i*_, *r*_*j*_) = (*v/L*)^2^*ij*Cov(*n*_*i*_, *n*_*j*_), where Cov(*n*_*i*_, *n*_*j*_) is obtained from Eq. (4d). In Figs. 8(a) and (b), we plot the mean and Fano factor of the local signal as a function of the gene segment *i*; note the contrast between the statistics of the fluorescent signal and the corresponding statistics of local RNAP – which is the statistics of nascent RNA – shown in Figs. 2(a) and (c).

**Figure 8:**
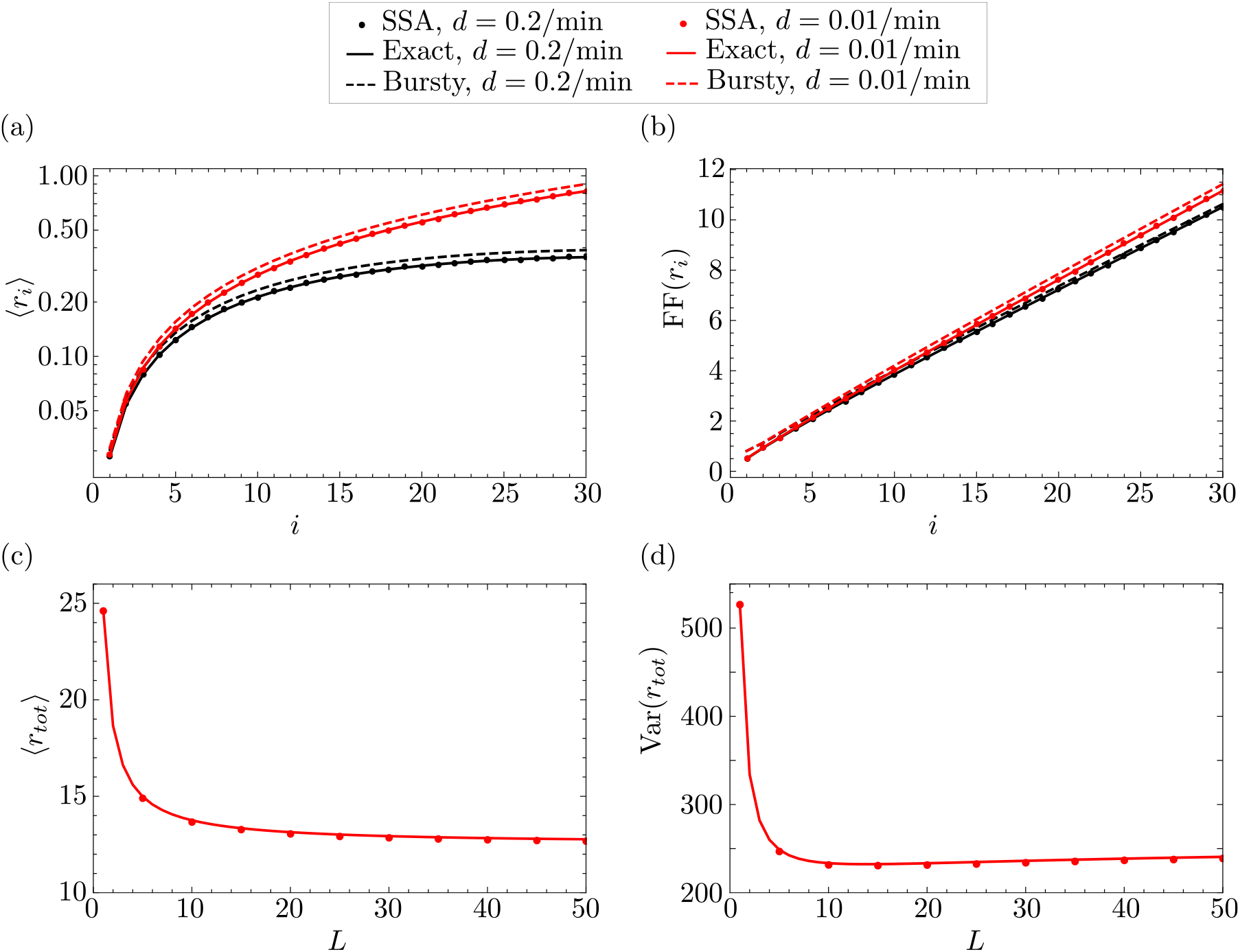
First and second moments of the local and total fluorescent signal for the bursty gene PDR5 in yeast. In panels (a) and (b), we show the dependence of the mean and the Fano factor of local fluorescent signal fluctuations on the gene segment *i*, as predicted by our exact theory (solid lines) and SSA (dots), respectively. The plots for *CV*^2^(*r*_*i*_) and *CC*(*r*_*i*_, *r*_*i*+1_) are identical to those of *CV*^2^(*n*_*i*_) and *CC*(*n*_*i*_, *n*_*i*+1_) in Fig. 2. The number of gene segments is arbitrarily chosen to be *L* = 30. In panels (c) and (d), we show the dependence of the mean and variance of total fluorescent signal fluctuations on the number of gene segments *L*, as predicted by our exact theory (Eq. (35); solid lines) and SSA (dots). The parameters *s*_*u*_, *s*_*b*_, *r*, and 〈*T*〉 are characteristic of the PDR5 gene and take the same values as in Fig. 2, as do the rates of elongation and RNAP detachment. The value of the parameter *v* is arbitrarily chosen to be *v* = 10.

Similarly, denoting by 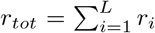 the total fluorescent signal across the gene, we find the following expressions for the steady-state mean 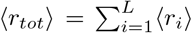 and the steady-state variance 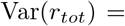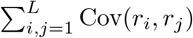:

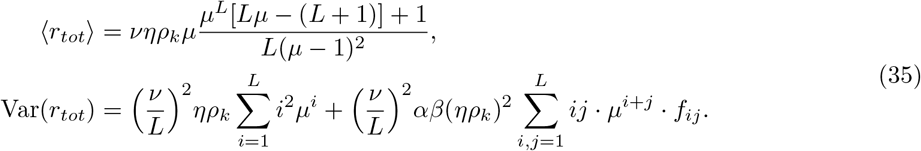

For a detailed derivation of the variance in Eq. (35), see Eq. (F.1) in Appendix F; see also Appendix G for the moments expressions in the bursty, constitutive, and deterministic elongation limits. In Figs. 8(c) and (d), we show the mean and the Fano factor of the total signal as a function of the number of gene segments (*L*); as above, note the contrasting difference between the statistics of the fluorescent signal and the corresponding statistics of total RNAP – which is the statistics of total nascent RNA – shown in Figs. 4(c) and (d).

Hence, the calculation of the statistics of the number of nascent RNAs from the raw signal intensity presents a challenge and has to be approached carefully. The expressions presented above allow for the inference of transcriptional parameters from the first two moments of the fluorescent signal by means of moment-based inference techniques [29]. Quantitative information about nascent RNA can also be obtained from electron micrograph images [30], which avoids the challenges presented by smFISH.

## 5 Model extension with pausing of RNAP

Thus far, we have studied a model where RNAPs do not pause as they move along the gene. A natural extension is provided by a modified model in which RNAPs pause along the gene at random sites and elongation is characterized by three processes: forward hopping, pausing, and unpausing of RNAP. The motivation for studying this extended model, which has recently been considered via stochastic simulation in [19], is that experiments have revealed that RNAP exhibits pauses of varying duration, typically on the timescale of few seconds [31,32].

### 5.1 Closed-form expressions for moments of local RNAP fluctuations

We extend the model described in Fig. 1 by assuming that the RNAP on gene segment *i* can switch between a non-paused (actively moving) state *P*_*i*_ and a paused state 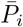. The actively moving state *P*_*i*_ switches to 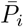 with rate *r*_*p*_, while the reverse reaction occurs with rate *r*_*a*_. Premature detachment from the actively moving RNAP occurs with rate *d*_*a*_, whereas it occurs with rate *d*_*p*_ from the paused RNAP. The resulting extended model is illustrated in Fig. 9(a). In Appendix A, we derive the mean and variance of the corresponding elongation time, which is not Erlang distributed now, as was the case for the model without pausing. Furthermore we find two interesting properties of the coefficient of variation 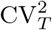 of the elongation time: (i) in the limit of large *L* at constant mean elongation time, 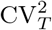 does not tend to zero, which implies that elongation is not deterministic; (ii) for small rates of premature detachment, 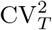 is at its maximum when *r*_*p*_ ≈ *r*_*a*_, i.e. when RNAP spends roughly half of its time in the paused state; see Appendix A for details and Fig. 9(b) for a confirmation through stochastic simulation.

**Figure 9:**
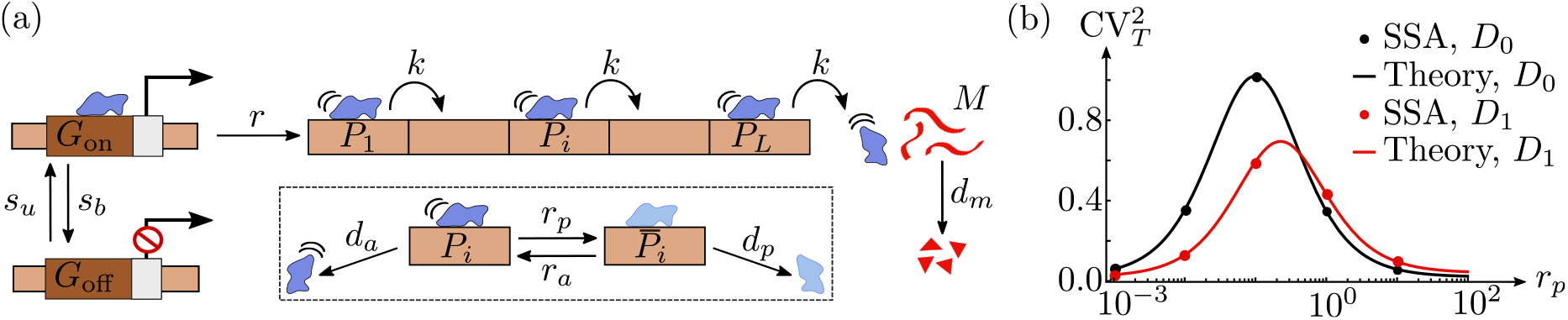
Model of transcription that includes RNAP pausing. In panel (a), we extend the model in Fig. 1 so that it takes into account pausing of RNAP at random segments on the gene. Pausing on gene segment *i* is modelled by the transition from the active state *P*_*i*_ to the paused state 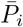 with rate *r*_*p*_, while the reverse (‘unpausing’) transition occurs with rate *r_a_*. Premature termination of RNAP occurs with rate *d*_*a*_ from the actively moving state, and with rate *d*_*p*_ from the paused state. In panel (*b*), we show the dependence of the coefficient of variation squared 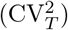 of the elongation time distribution on the pausing rate (*r*_*p*_), as predicted from SSA (dots) and theory (Eq. (A.7); solid lines). Results are shown for two different parameter regimes: *D*_0_ ≡ {*d*_*a*_ = 0/min = *d*_*p*_} (no premature polymerase detachment) and *D*_1_ ≡ {*d*_*a*_ = 0.05/min = *d*_*p*_} (premature polymerase detachment). The remaining parameters are fixed to *L* = 50, *k* = 10/min, and *r*_*a*_ = 0.1/min.

#### Proposition 3.

*Let the number of RNAP molecules in the active state P_i_ be denoted by* 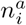, *let the number of molecules in the paused state* 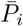 *be* 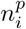, *and let the number of molecules of mature RNA be denoted by n. Let σ* = *r_p_/r_a_ be the ratio of the pausing and activation rates, let* 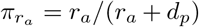 *be the probability of RNAP switching to the actively moving state from the paused state, and let* 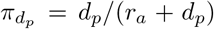 *be the probability of premature RNAP detachment from the paused state. Furthermore, we define the new parameters* 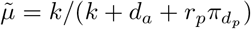 *and* 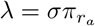.

*Then, it follows that the steady-state mean number of RNAP molecules in the active and paused states on gene segment i (i* = 1, … *L) is given by*

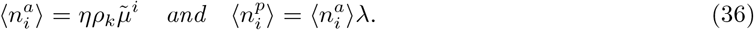

*Hence, the total mean number of RNAP molecules on each gene segment i reads*

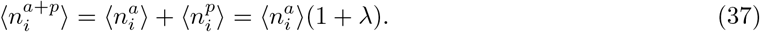

The proof of Prop. 3 can be found in Appendix H. Note that in the limit of no pausing, i.e. for *r*_*p*_ = 0, Eq. (37) reduces to the expression for the mean of RNAP reported in Eq. (2b).

#### Proposition 4.

*Let* 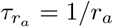 *be the timescale of RNAP activation from the paused state, let* 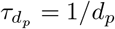 *be the timescale of premature termination of paused RNAP, let τ_p_* = 1/(*k* + *d*_*a*_) *be the typical time that an actively moving RNAP spends on a gene segment, and let τ_pp_* = 1/(*r*_*a*_ + *d*_*p*_) *be the typical time spent in the paused state. Additionally, we define new parameters* 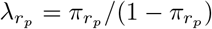, *where* 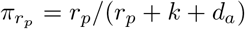 *is the probability of the actively moving RNAP switching to the paused state, as well as*

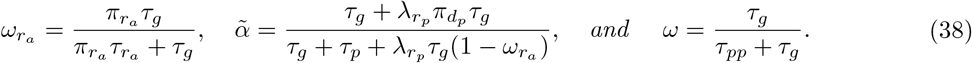

*Assume that the elongation rate is faster than the rates of RNAP pausing, activation, and premature termination, i.e. that k* ≫ *r_a_, r_p_, d_a_, d_p_. Then, it follows that to leading order in* 1/*k, asymptotic expressions for the variances and covariances of molecule number fluctuations of active and paused RNAP are given by*

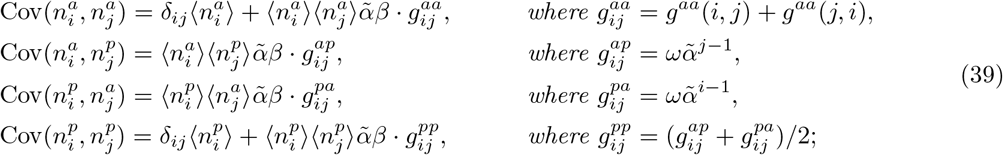

*here, i, j* = 1, 2, …, *L and*

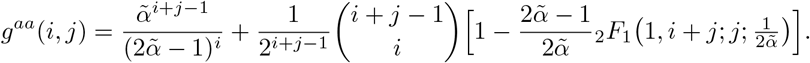

These results are proved in full in Appendix H. From Appendix A, we also have that the mean elongation time in the pausing model is given by

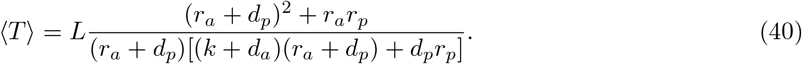

Solving Eq. (40) for the elongation rate *k*, we find that in the limit of *L* → ∞ taken at constant mean elongation time, *k* tends to infinity and hence is much larger than *r*_*a*_, *r*_*p*_, *d*_*a*_, and *d*_*p*_, which implies that the results of Prop. 4 hold naturally in that limit.

### 5.2 Approximate distributions of total RNAP and mature RNA

#### Negative Binomial approximation of total RNAP distribution

We define the total number of RNAP molecules as 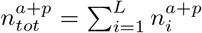. It then immediately follows from Eq. (37) that the mean of the total RNAP distribution in the pausing model is given by

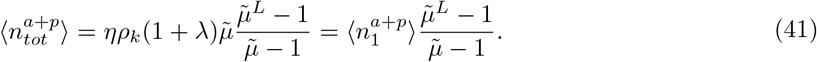

It can also be shown that the variance of total RNAP fluctuations reads

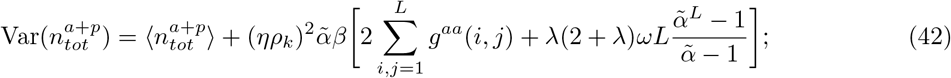

see Appendix H. Next, we approximate the distribution of total RNAP by a Negative Binomial distribution whose mean and variance match those just derived, i.e. we use Eq. (20) with 〈*n*_*tot*_〉 changed to 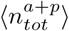, and Var(*n*_*tot*_) replaced by 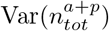. The resulting approximate Negative Binomial distribution is compared with the distribtution obtained from SSA in Figs. 10(a) and (b) for two different yeast genes, PDR5 and DOA1. The results verify that our approximation is accurate provided the elongation rate *k* is significantly larger than the other parameters, as assumed in Prop. 4.

**Figure 10:**
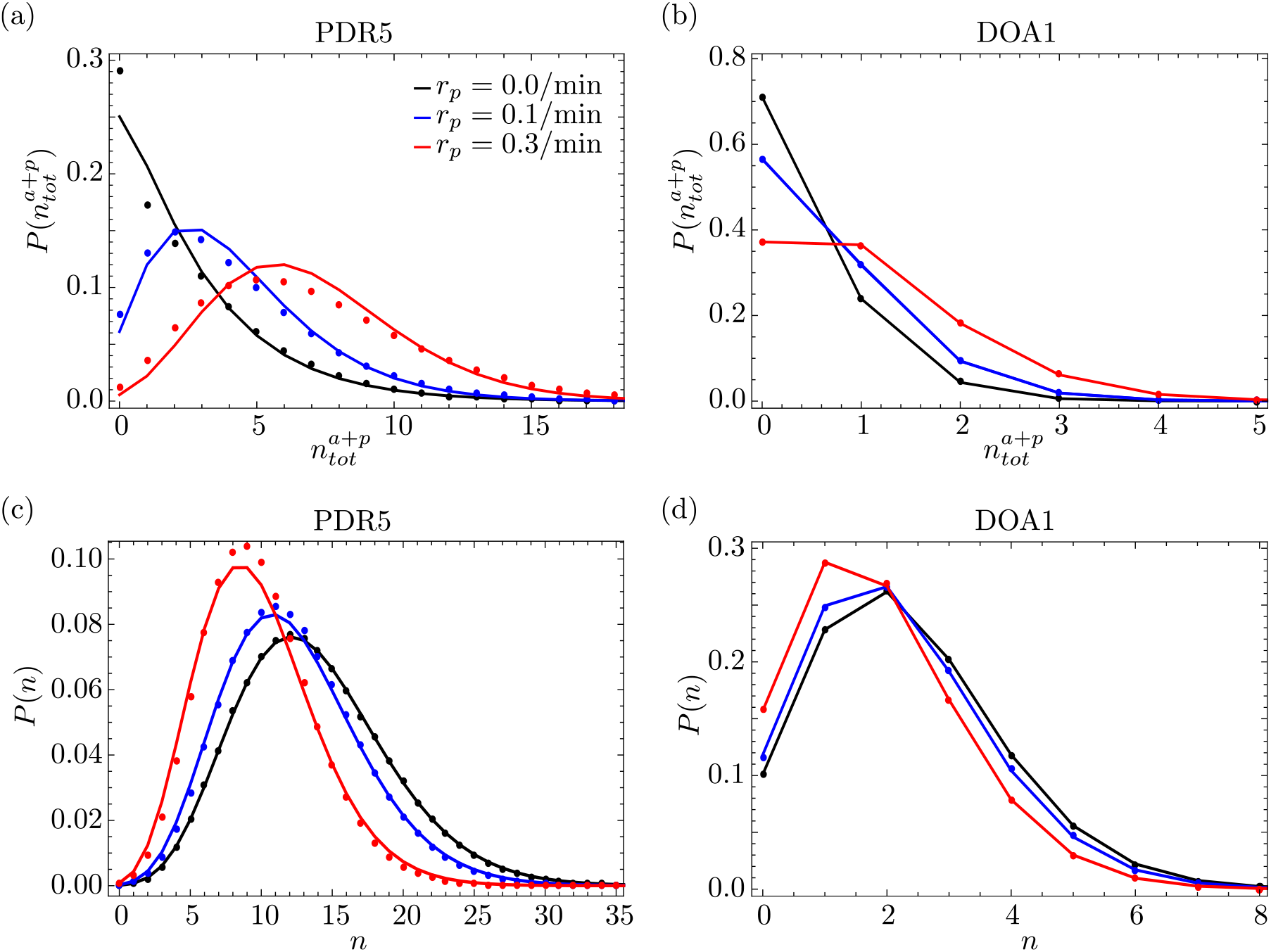
Dependence of the steady-state probability distributions of total RNAP and mature RNA on the RNAP pausing rate *r*_*p*_ for two different genes in yeast. In panels (a) and (b), we compare the distribution 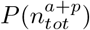 of the total number of RNAP molecules, as predicted by our model (solid lines), with that obtained from SSA (dots) for yeast genes PDR5 and DOA1, respectively. The model prediction involves fitting a Negative Binomial distribution with a mean and variance that are given by the closed-form expressions in Eqs. (41) and (42). In panels (c) and (d), we compare the distribution *P* (*n*) of mature RNA, as obtained from singular perturbation theory (Eq. (43); solid lines) with the SSA (dots) for yeast genes PDR5 and DOA1, respectively. Note that for both genes, we keep all parameters fixed (including the elongation rate *k*) while varying the pausing rate *r*_*p*_ to simulate an experiment where the pausing rate can be perturbed directly. The parameters for each gene can be found in Table 1a; we furthermore used *L* = 50 and fixed *k* to *L* /〈*T*〉, where 〈*T*〉 is the mean elongation time measured experimentally and reported in Table 1a. Note that the actual mean elongation time is not fixed, as it depends on the pausing rate (*r*_*p*_) via Eq. (40). The remaining parameters are fixed to *r*_*a*_ = 0.1/min, *d*_*a*_ = 0.01/min, and *d*_*p*_ = 0.03/min. The value of *d*_*a*_ is taken from Table 1 in [17], where it is reported as the premature termination rate of polymerase in *E. coli*; the value of *d*_*p*_ was chosen to be larger than that of *d*_*a*_ to simulate a scenario where premature detachment is enhanced in the paused state. Note that our theory is less accurate for PDR5 than it is for DOA1, as all parameters are very small compared to the elongation rate in the latter case, hence satisfying better the assumptions behind the theory.

#### Perturbative approximation of mature RNA distribution

We can apply singular perturbation theory to formally derive the distribution of mature RNA, assuming that *k*/*d*_*m*_ ≫ 1 and *r*_*a*_/*d*_*m*_ ≫ 1. Following the derivation in Section 3.2, we find the following analytical expression for the steady-state probability distribution of mature RNA:

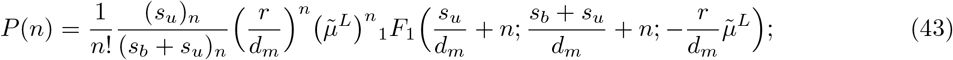

see Appendix I for details. Note that the solution in Eq. (43) is dependent on the parameter 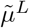, which gives the probability that an RNAP molecule does not prematurely detach before termination; see Appendix A. Also, note that in the limit of zero premature termination, i.e. for *d*_*a*_ = 0 = *d*_*p*_, Eq. (43) is identical to the distribution of mature RNA predicted by the telegraph model. Finally, by solving Eq. (40) for *k*, then substituting the resulting expression in Eq. (43) and taking the long-gene limit of *L* → ∞ at constant 〈*T*〉, we obtain that the probability distribution of mature RNA has the same functional form as in Eq. (43), but with

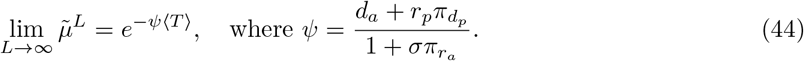

Note that Eq. (43) with Eq. (44) equals the steady-state solution predicted by the telegraph model, with the initiation rate *r* renormalised to *re*^−*ψ*〈*T*〉^. In Figs. 10 (c) and (d), we verify the accuracy of our analytical solution using stochastic simulation for two different genes in yeast. Note that the effect of changing the pausing rate *r*_*p*_ has relatively little effect on the distribution of mature RNA, as compared to the effect on the distribution of total RNAP; cf. panels (a) and (b) of Fig. 10 in comparison with panels (c) and (d), respectively.

## 6 Summary and Conclusion

In this paper, we have analyzed a detailed stochastic model of transcription. Our model extends previous analytical work [7, 9] by (i) taking into account salient processes, such as premature detachment and pausing of RNAP, that were previously not considered analytically; (ii) deriving explicit expressions for the mean and variance of RNAP numbers (nascent RNA) on gene segments as well as on the entire gene; deriving explicit expressions for the mean and variance of the fluorescent nascent RNA signal obtained from smFISH, and identifying differences between the statistics thereof and those of direct measurements of nascent RNA; (iv) finding approximate distributions of total nascent RNA fluctuations on a gene, without assuming slow promoter switching. A number of interesting observations from our work include the following.

i. When the premature detachment rate of RNAP is non-zero and gene expression is bursty, the coefficient of variation of local RNAP fluctuations can either decrease or increase with distance from the promoter. By contrast, when expression is constitutive, the coefficient of variation increases monotonically with distance from the promoter. Other statistical measures such as the mean, Fano factor, and correlation coefficient of local RNAP numbers decrease monotonically with distance from the promoter.
ii. In the limits of bursty expression, deterministic elongation, and no premature detachment or pausing, the Fano factor of total nascent RNA equals 1 + 2*b*, whereas that of mature RNA is 1 + *b*, where *b* denotes the mean burst size. An implication is that using the telegraph model to estimate the mean burst size from nascent RNA data will result in an overestimate by a factor of 2. Another implication is that deviations from Poisson fluctuations are more apparent in data for nascent RNA than they are for mature RNA. One can further state the following relationship: the Fano factor of nascent RNA equals twice the Fano factor of mature RNA, minus 1. If expression is non-bursty, then the Fano factor of nascent RNA can be larger or smaller than that of mature RNA, as determined by the condition in Eq. (19).
iii. For genes characterized by bursty expression, the sensitivity of the noise in total RNAP fluctuations is highest to perturbations in the gene activation rate and the mean elongation time; for constitutive genes, the most sensitive parameters are the initiation rate and the mean elongation time.
iv. A Negative Binomial distribution, parameterized with the expressions for the mean and variance of total nascent RNA derived here, provides a good approximation to the true distribution of total nascent RNA fluctuations on a gene when expression is either bursty or constitutive; the approximation is not accurate when the gene spends roughly equal amounts of time in the active and inactive states. We show that the distribution of nascent RNA is highly sensitive to the distribution of elongation times. In particular, if the elongation time is assumed to be exponentially distributed, as is implicitly assumed by telegraph models of nascent RNA, then the probability of observing zero RNA is much lower than if the elongation time is assumed to be fixed.
v. Using geometric singular perturbation theory (GSPT), we have rigorously proven that, in the limit of deterministic elongation (or fast elongation), no pausing and premature detachment, the steady-state distribution solution of mature RNA in our model is identical to that in the telegraph model [25]. Consideration of pausing and premature detachment leads to a distribution that can also be obtained from a telegraph model with appropriately renormalized parameters.

The main limiting assumption of our theoretical approach is that the initiation rate is slow enough such that RNAP molecules do not collide with each other while moving along the gene. Hence, the expressions we have derived are reasonable for all but the strongest promoters which are characterized by very fast initiation rates. We anticipate that approximate closed-form expressions for the corresponding moments can also be derived when volume exclusion between RNAPs is taken into account by a modification of methods previously devised to understand molecular movement and kinetics in crowded conditions [33,34]. It is also possible to extend our model by including translation of mature RNA to protein; one can then again apply GSPT to derive distributions for protein numbers in the limit of RNA decaying much faster than protein; however, given item (v) above, we anticipate that the resulting protein distribution will be very similar to those derived from models that do not explicitly take into account nascent RNA [35, 36]. Further research is required to develop simple approximations of the nascent RNA distribution that are accurate independent of the ratio of gene switching rates. Finally, given the strong recent interest in the development of statistical inference techniques in molecular biology [29, 37, 38], we expect that our closed-form expressions for the moments and distributions of nascent and mature RNA will be useful for developing computationally efficient and accurate methods for estimating transcriptional parameters.

## Acknowledgments

R.G. acknowledges useful discussions with Zhixing Cao and Tineke Lenkstra. This work was supported by a departmental PhD studentship to T.F.

## Appendix

## A Distribution of elongation time

In this section, we answer the following question: what is the distribution of the elongation time, i.e. the time between initiation and termination? In other words, with reference to Fig. 9 – which includes the non-pausing model in Fig. 1 as a special case – we want to find the distribution of the time at which RNAP leaves gene segment *L* (termination) if it was in the active state on gene segment 1 at *t* = 0 (initiation).

Let *z*_*i*_(*t*) be the probability of an RNAP to be on gene segment *i* in the active state at time *t*, let 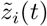 be the probability of the RNAP to be on gene segment *i* in the paused state at time *t*, and let 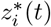 be the probability of the RNAP moving to gene segment *i* + 1 at time *t*; note that 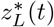 is the probability of the RNAP falling off the gene and forming a mature RNA, since for *i* = *L*, gene segment *L* + 1 does not exist. Then, it follows from the reaction scheme illustrated in Fig. 9 that the master equations describing the Markovian dynamics on gene segment *i* are given by:

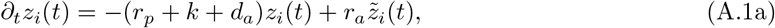

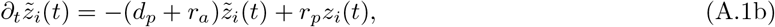

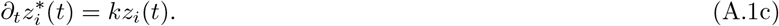

Now, we use these equations to find the distribution of the time when RNAP jumps to gene segment *i* + 1, given that it is on gene segment *i* in the active state at *t* = 0, i.e. that *z*_*i*_(0) = 1 and 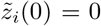. Taking the Laplace transform of Eqs. (A.1a) and (A.1b), we find

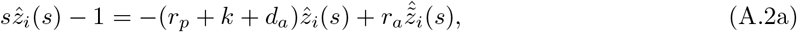

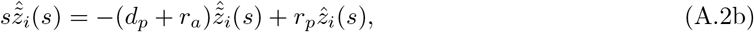

where 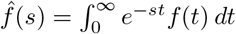. Solving these equations simultaneously, we obtain

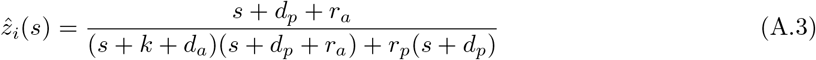

Let *w*(*t*)*dt* be the probability that the RNAP moves from segment *i* to *i* + 1 in the time interval (*t, t* + *dt*). Then, it follows from Eq. (A.1c) that 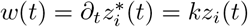. Integrating *w*(*t*) over all times gives us the probability that the RNAP ultimately moves to the next segment *i* + 1

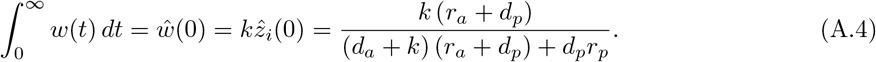

Note that 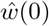 is identical to the parameter 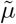, as defined in Prop. 3. Let *y*(*t*)*dt* be the probability that the RNAP moves from gene segment *i* to segement *i*+1 in the time interval (*t, t* + *dt*), *conditioned* on those realisations that lead to an RNAP moving to the next gene segment *i* + 1. (In other wordds, we exclude those realisations that lead to premature detachment.) Then, it follows by the definition of conditional probabilities that 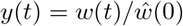, which implies

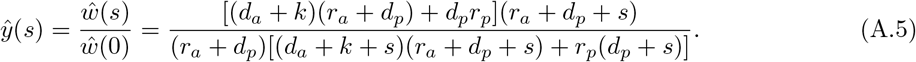

It follows that the mean 〈*t*〉 and variance Var(*t*) of the time *t* it takes RNAP to move to the next gene segment are given by

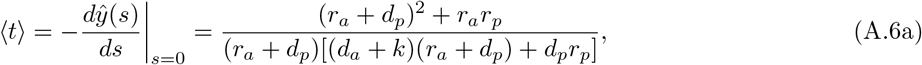

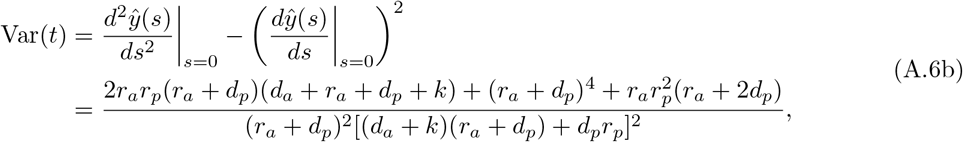

respectively. Since RNAP can only move forwards in our model (irreversible motion), it follows that the time it takes an RNAP to move from the *i*-th gene segment to the (*i* + 1)-th is independent of the time taken to move from another, *j*-th segment *j* to the (*j* + 1)-th. Hence, the time required for an RNAP to move across the entire gene from the first to the *L*-th segment, i.e. the time *T* from initiation to termination (the elongation time), is a sum of *L* independent and identical random variables. Thus, we can immediately state that the mean elongation time is 〈*T*〉 = *L*〈*t*〉, whereas the variance of the elongation time is Var(*T*) = *L*Var(*t*). The coefficient of variation squared takes the form

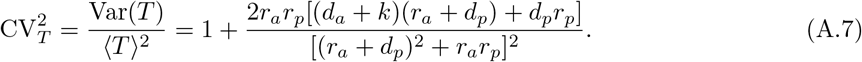

From Eq. (A.7), it can be shown that for small premature detachment rates, the coefficient of variation of the elongation time is maximized when *r*_*p*_ ≈ *r*_*a*_. Taking the limit of infinitely many gene segments at constant mean elongation time, i.e. solving for *k* from the expression for the mean elongation time in Eq. (A.6), substituting into Eq. (A.7), and taking the limit of *L* → ∞, we obtain

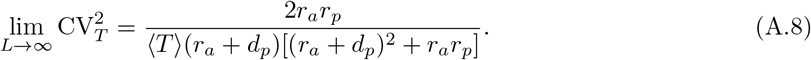

For the non-pausing model shown in Fig. 1, the above results simplify considerably due to *r*_*p*_ = 0 = *d*_*p*_ and *d*_*a*_ = *d*; in that case, the inverse Laplace transform of Eq. (A.5) implies that *y*(*t*) is an exponential distribution with parameter *k* + *d*. Hence, the total time it takes an RNAP to move across the entire the gene is the sum of *L* independent and identically distributed exponential random variables, i.e. an Erlang distribution with shape parameter *L* and rate *k* + *d*, which implies that the mean elongation time is *L/*(*k* + *d*), with coefficient of variation 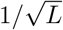. It can be seen from Eq. (A.8) that deterministic elongation can only be observed when there is no pausing, i.e. when *r*_*p*_ = 0.

## B Solution of Lyapunov equation

*Proof of Proposition 2.* We start by defining the symmetric functions *f*_*ij*_ = *f*_*ji*_ for *i, j* = 1, …, *L* as

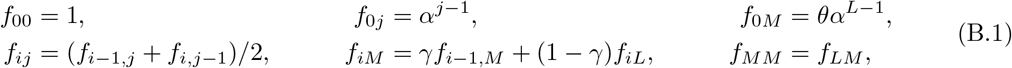

where the non-dimensional parameters *α*, *γ*, and *θ* are defined in Prop. 2. The elements of the Lyapunov equation given by Eq. (5) can be written explicitly as a set of simultaneous equations:

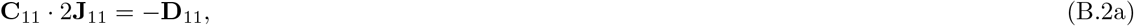

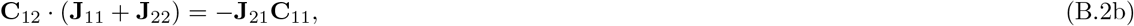

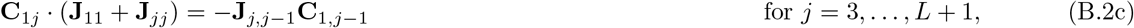

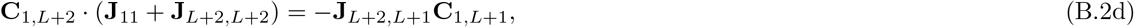

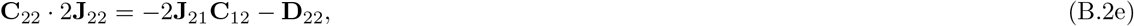

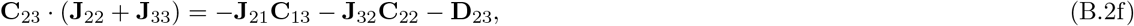

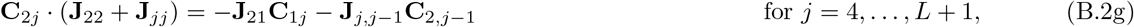

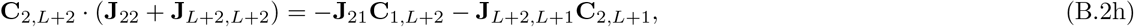

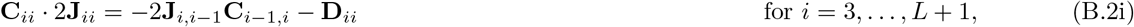

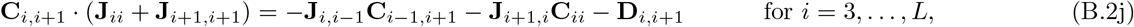

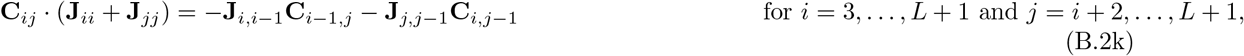

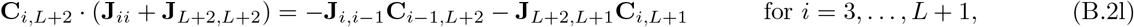

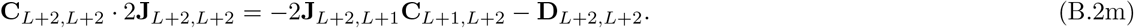

Now, we substitute the elements of the Jacobian matrix **J** and the diffusion matrix **D** from Eqs. (6) and (7), respectively, into the above system of algebraic equations, which we then solve to find the elements of the covariance matrix **C**. Note that, for the following mathematical derivation, we take into account the expressions for the steady-state mean numbers of species given in Eq. (2), as well as the definition of the functions *f*_*ij*_ in Eq. (B.1).

From Eq. (B.2a), one easily obtains **C**_11_ = *η*^2^*β*. Then, it follows from Eq. (B.2b) that

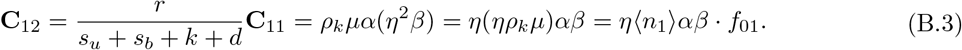

Eq. (B.2c) implies that, for *j* = 3, …, *L* + 1:

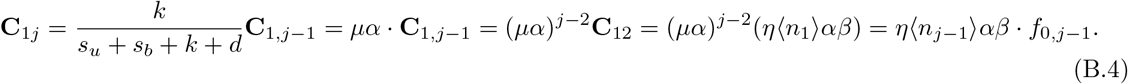

From Eq. (B.2d), we have that

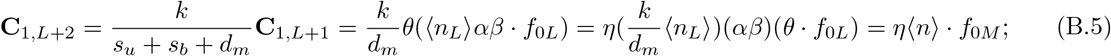

from Eq. (B.2e), we find

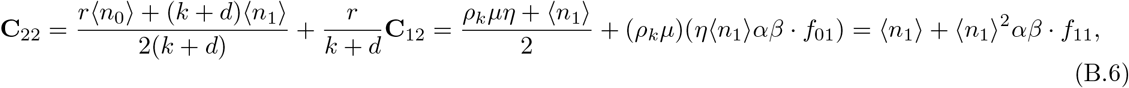

since *f*_11_ = (*f*_01_ + *f*_10_)/2 = *f*_01_ from the definition in Eq. (B.1).

From Eq. (B.2f), we obtain

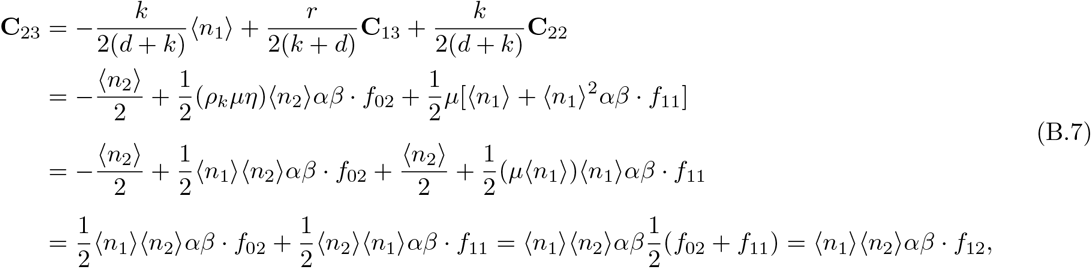

since *f*_12_ = (*f*_02_ + *f*_11_)/2 from the definition in Eq. (B.1).

From Eq. (B.2g), we have that, for *j* = 4, …, *L* + 1,

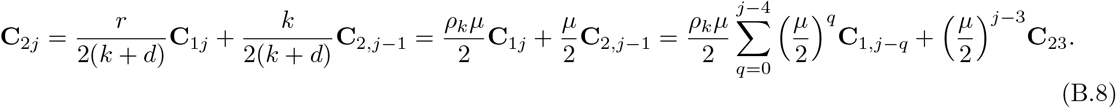

The proof of Eq. (B.8) is given in Lemma B.1. The above expression for **C**_2*j*_ can be further simplified to

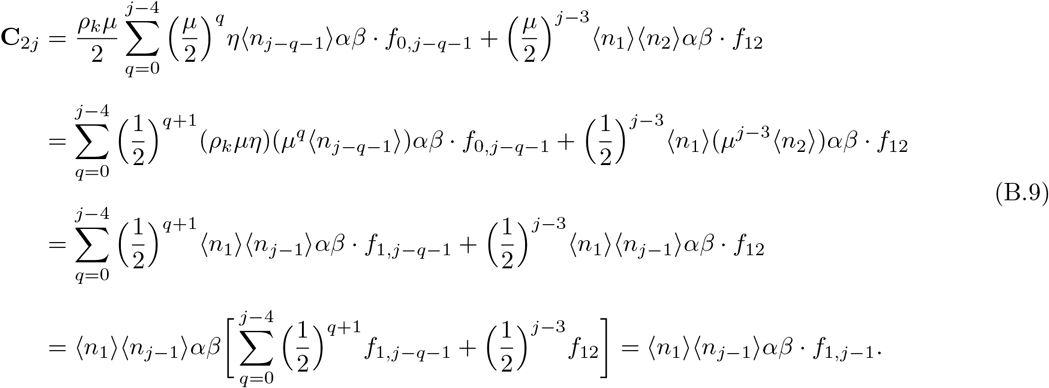

For the proof of the last equality in Eq. (B.9), see Lemma B.2.

From Eq. (B.2h), we have that

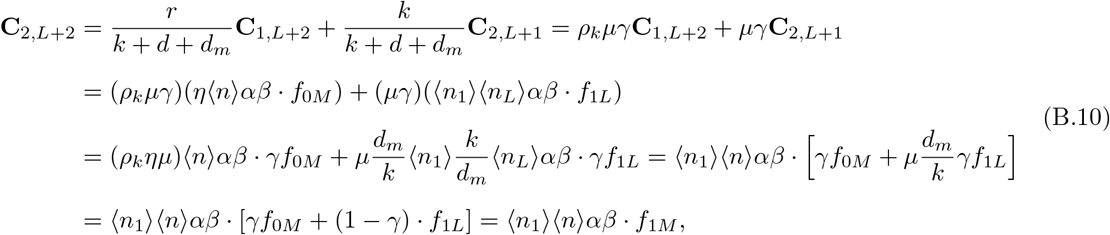

where *f*_1*M*_ is defined in Eq. (B.1).

Eqs. (B.2i) through (B.2k) yield the system

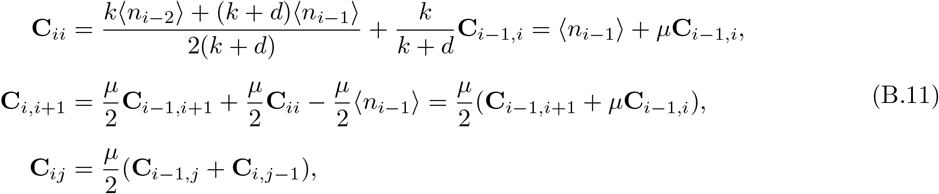

which can be rewritten more compactly as

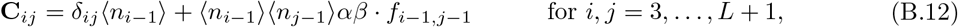

where *δ*_*ij*_ is the Kronecker delta. A detailed derivation is given in Lemma B.3.

From Eq. (B.2l), we have that for *i* = 3, …, *L* + 1,

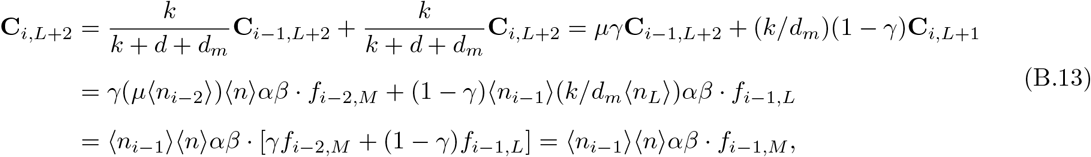

where *f*_*iM*_ is defined in Eq. (B.1).

Finally, Eq. (B.2m) yields

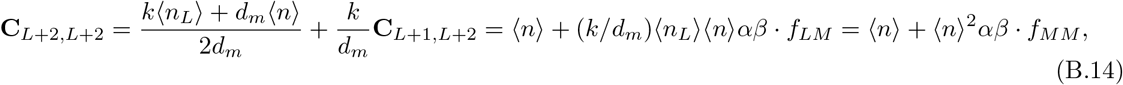

where *f*_*MM*_ = *f*_*LM*_ is defined in Eq. (B.1).

Summarizing the above results, we conclude that the solution for the symmetric covariance matrix **C** is given by the system in Eq. (4), where we have that Cov(*n*_*i*_, *n*_*j*_) = **C**_*i*+1*,j*+1_, Cov(*n*_*i*_, *n*) = **C**_*i*+1,*L*+2_ for *i, j* = 0, …, *L* and Var(*n, n*) = **C**_*L*+2,*L*+2_. Here, the functions *f*_*ij*_ are defined as in Eq. (B.1). Now, the recurrence relation *f*_*ij*_ = (*f*_*i*−1*,j*_ + *f*_*i,j*−1_)/2 in Eq. (B.1) can be solved for *i, j* = 1, 2, …, *L* via the method of generating functions, which gives the following analytical expression:

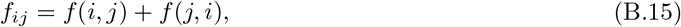

where

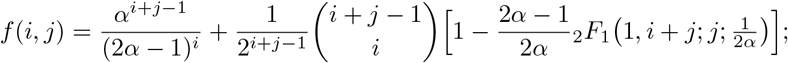

see Lemma B.5 for a detailed derivation. Additionally, we can easily prove that the function *f*_*iM*_ in Eq. (B.1) can be rewritten as

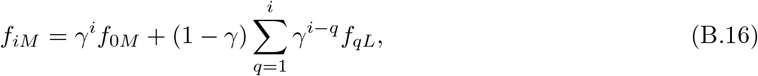

as shown in Lemma B.4.

### Lemma B.1.

*For j* = 4, …, *L* + 1, *we have the identity*

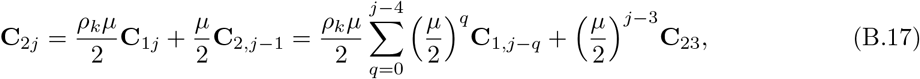

*as stated in Eq. (B.8).*

*Proof.* The identity in Eq. (B.17) will be proved by induction: one can easily show that it holds for *j* = 4.

Now, we assume that Eq. (B.17) is true for some *j* ≥ 5; hence, for *j* + 1, we have

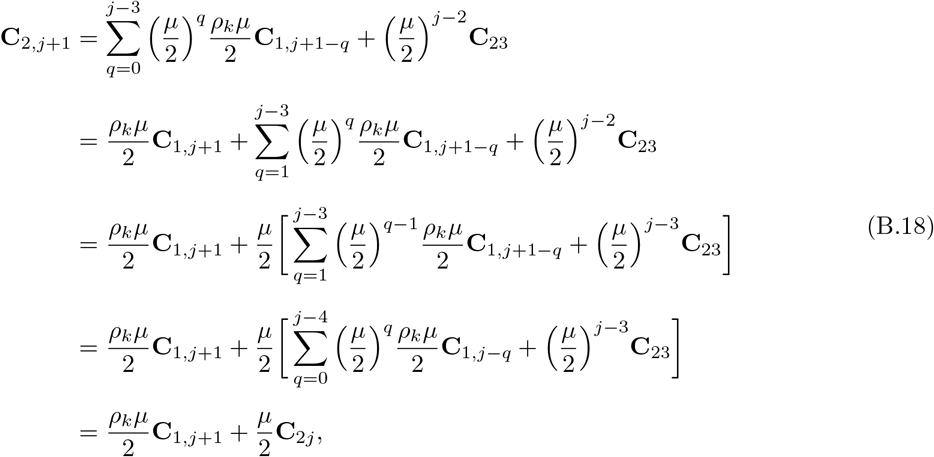

as claimed, which implies that the identity in Eq. (B.17) holds for all *j* = 4, …, *L* + 1.

### Lemma B.2.

*The function f*_1*j*_, *which is defined by the recurrence relation f*_1*j*_ = (*f*_0*j*_ + *f*_1,*j*−1_)/2 *in Eq. (B.1), satisfies the identity*

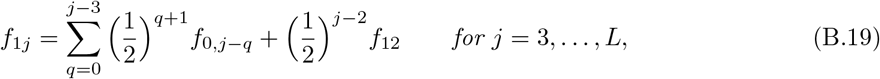

*as stated in Eq. (B.9).*

*Proof.* We will again prove Eq. (B.19) by induction. For *j* = 3, we have from Eq. (B.19) that *f*_13_ = (*f*_03_ + *f*_12_)/2, which is true by the definition of *f*_13_. We assume that the identity in Eq. (B.19) is correct for some *j ≥* 4; then, for *j* + 1, the definition of *f*_1,*j*+1_, in combination with our assumption, implies

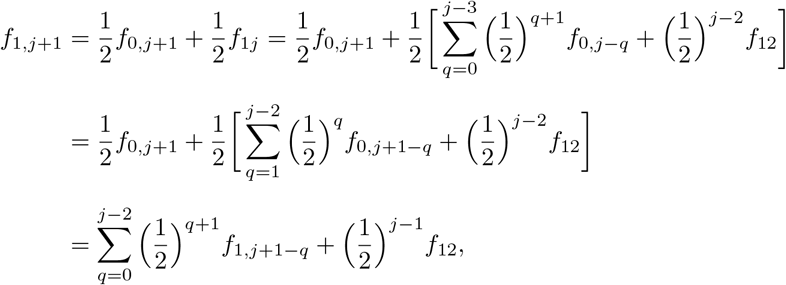

as claimed. Hence, the equality in Eq. (B.19) is true for all *j* = 3, …, *L*.

### Lemma B.3.

*The system in Eq. (B.11), which is given by*

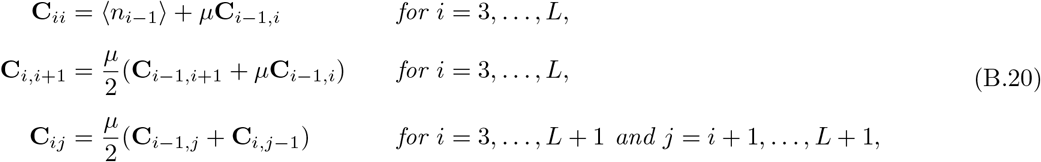

*is equivalent to the system*

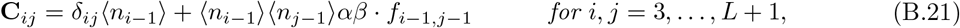

*as stated in Eq. (B.12). Here, the functions f_ij_ are defined as in Eq. (B.1).*

*Proof.* We again use the method of induction. For *i* = 3, we have

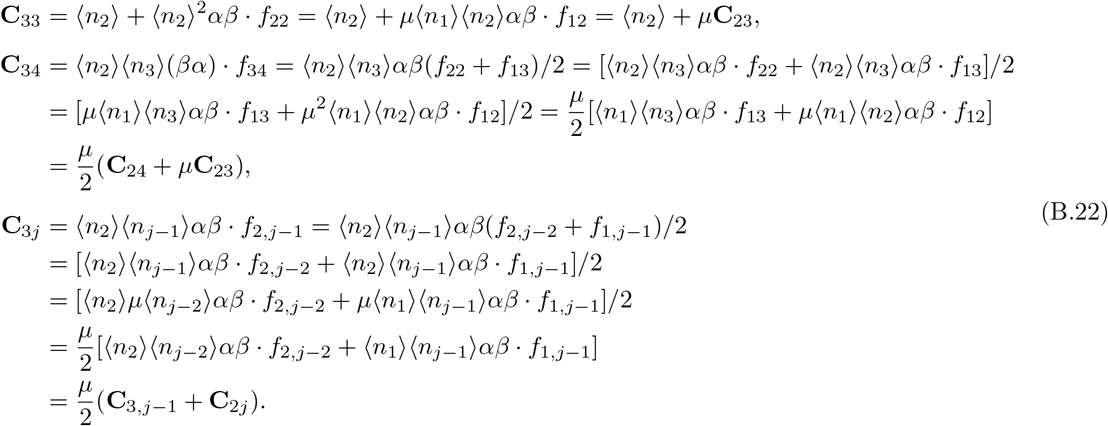

Now, we assume that the statement is true for some *i* ≥ 4; then, for *i* + 1, we have

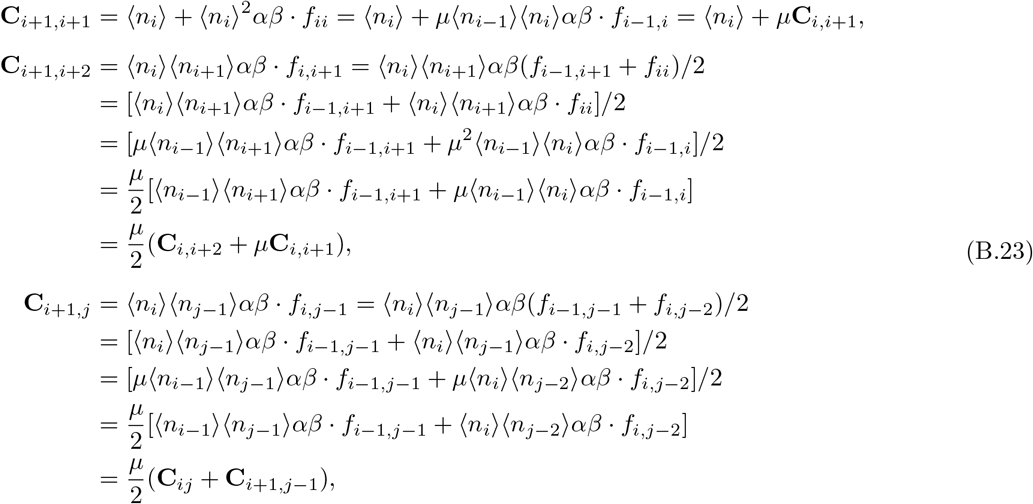

which is also correct. Hence, the statement of the lemma is true for all *i* and *j*, as stated.

### Lemma B.4.

*For i* = 1, …, *L, the function f_iM_ defined in Eq. (B.1) can be simplified as in Eq. (B.16); specifically, we have the identity*

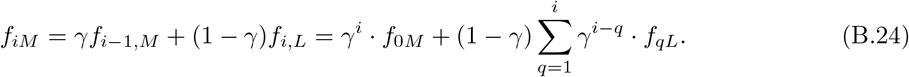

*Proof.* The proof is by induction: for *i* = 1, the identity is obvious. We now suppose that Eq. (B.24) is true for some *i* ≥ 2; hence, for *i* + 1, we have

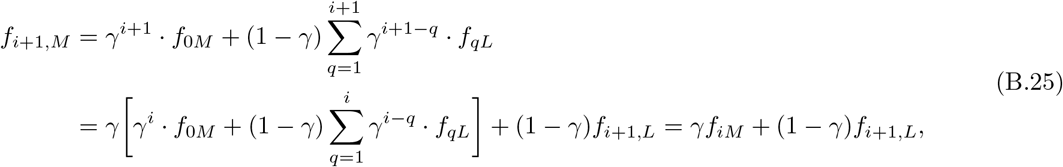

which is correct. Hence, Eq. (B.24) is true for all *i*, as stated.

### Lemma B.5.

*For i, j* = 1, …, *L, the solution of the recurrence relation f_ij_* = (*f*_*i,j*−1_ + *f*_*i*−1,*j*_)/2 *in Eq. (B.1) is given by f_ij_* = *f* (*i, j*) + *f* (*j, i*), *where*

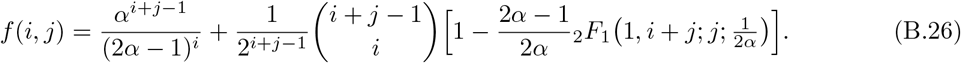

*Proof.* In order to solve the recurrence relation for the function *f*_*ij*_, we take into account the initial conditions *f*_00_ = 1 and *f*_0*j*_ = *f*_*j*0_ = *α*^*j*−1^. Then, we define a generating function *g*(*x, y*) via

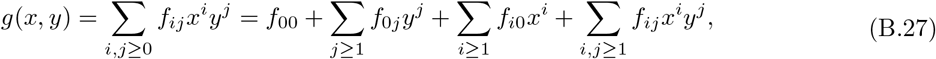

where the last term can be rewritten as

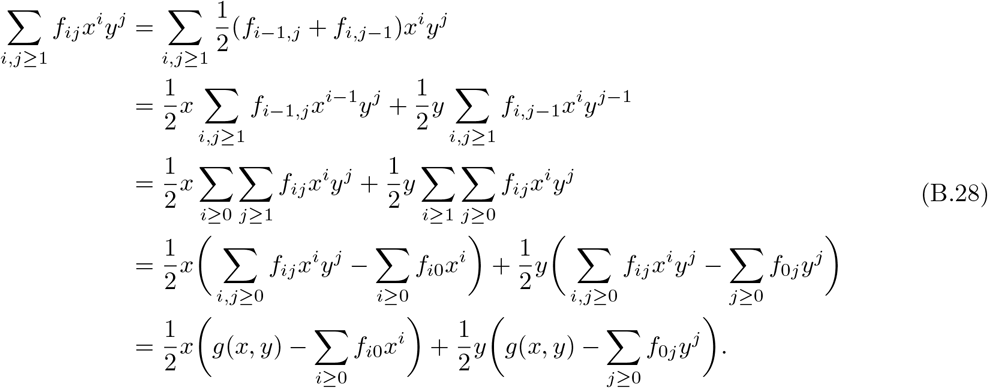

Hence, Eq. (B.27) becomes

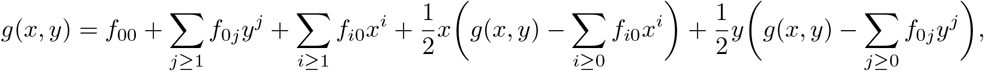

which is equivalent to

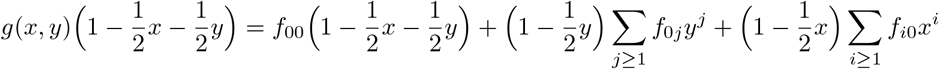

or

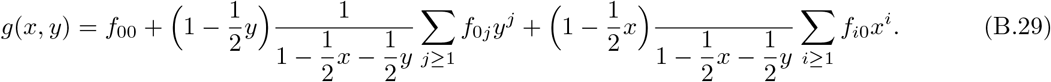

Taking into account the initial conditions it follows that

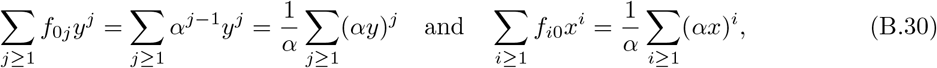

which we substitute into Eq. (B.29) to obtain

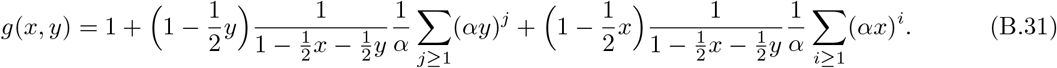

Making use of the well-known symmetric, bivariate generating function of the binomial coefficients

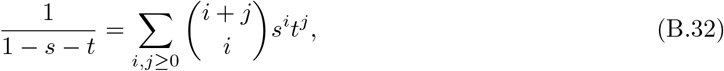

we can rewrite Eq. (B.31) as

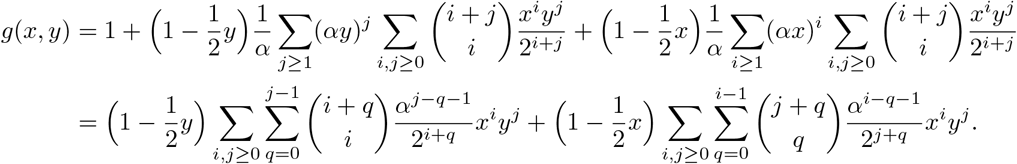

Rearranging sums in the above expression, we find

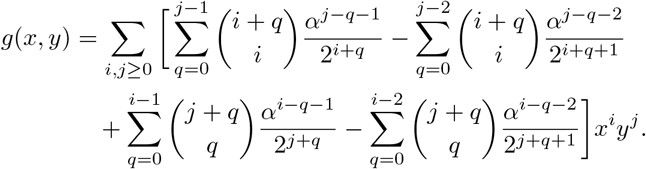

Hence, we obtain the following exact expression for the function *f*_*ij*_,

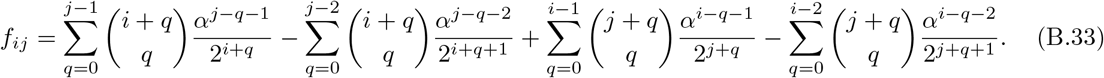

The expression in Eq. (B.33) can be simplified further due to its symmetry with respect to the indices *i* and *j*: we write *f*_*ij*_ = *f* (*i, j*) + *f* (*j, i*), where *f* (*i, j*) is defined as

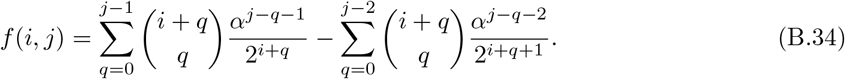

The function *f* (*i, j*) can be further simplified as

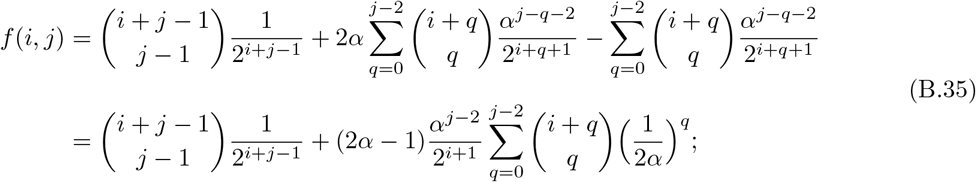

next, we use the identity

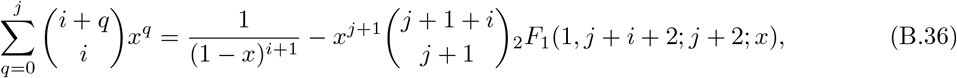

where _2_*F*_1_ is again the generalised hypergeometric function of the second kind [21]. Hence, Eq. (B.35) becomes

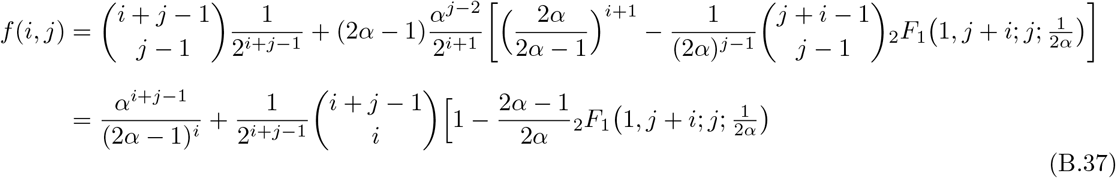

Given the expression for *f* (*i, j*) in Eq. (B.37), one can find the corresponding expression for *f* (*j, i*) by exchanging the indexes *i* ↔ *j*.

## C Variance of total RNAP distribution

In this section, we derive the exact expression for the variance of the total RNAP distribution, as stated in Eq. (10), which is given by the sum over the covariances Cov(*x*_*i*_, *x*_*j*_) (*i, j* ∈ {1, …, *L*}), as defined in Eq. (4d). Hence, we have

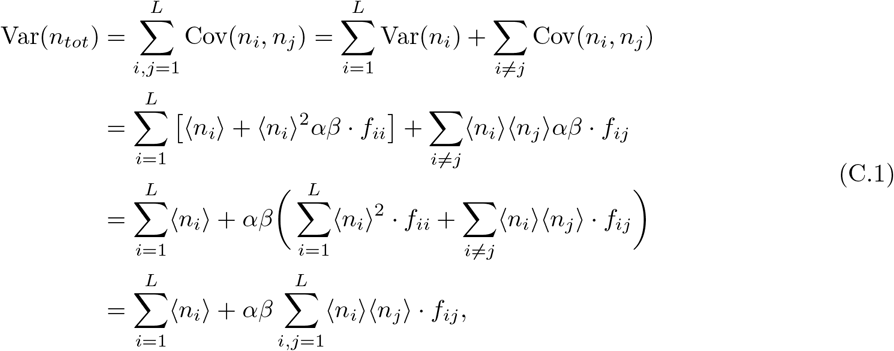

where the function 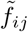 is given in Eq. (10). The first term in Eq. (C.1) equals 〈*n*_*tot*_〉, the mean of the total RNAP distribution, as given in Eq. (10); substituting in the expressions for the means 〈*n*_*i*_〉 from Eq. (2b), as well, we obtain

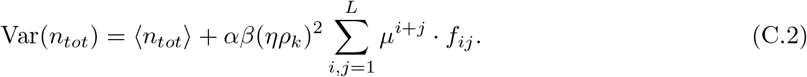

### Lemma C.1.

*In the limit of deterministic elongation, i.e. for* L → ∞ *the expression for* Var(*n*_*tot*_) *in Eq. (10) simplifies to*

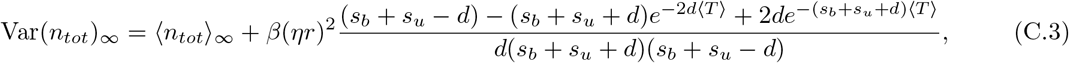

*which can be further simplified to the expression in Eq. (11).*

*Proof.* In order to find the limit of *L* → ∞ in Eq. (10) (or Eq. (C.2)), we have to evaluate the term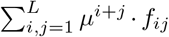 in that limit. For the following derivation, we consider the function *f*_*ij*_ = *f* (*i, j*) + *f* (*j, i*), where *f* (*i, j*) is defined in in terms of sums in Eq. (B.34). Hence, we have

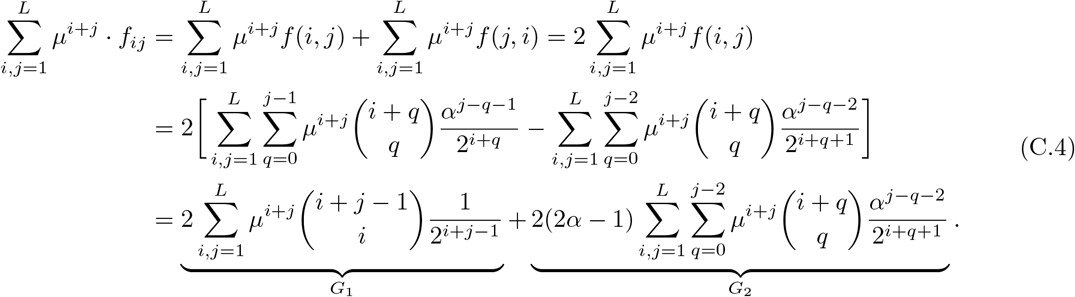

Substituting *k* → *L*/〈*T*〉 − *d* in Eq. (C.4) and taking the limit of *L* → ∞, we have that 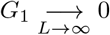; hence, Var(*n*_*tot*_) evaluates to

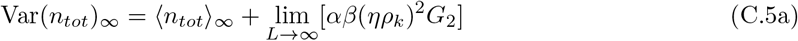

in that limit, which yields the expression in Eq. (C.3), as can easily be verified with the computer algebra package Mathematica. Hence, in the limit of deterministic elongation, the expression for the variance of the RNAP distribution in Eq. (10) reduces to the one in Eq. (11), as claimed.

## D Moments of total RNAP and mature RNA in bursty and constitutive limits

## Moments of total RNAP in the bursty limit

In the bursty limit, the expressions for the mean and variance of the total RNAP distribution given in Eq. (10) simplify to

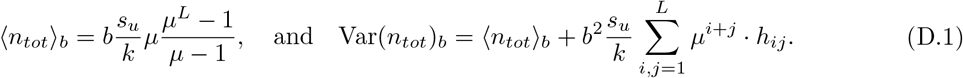

If, furthermore, we take the limit of deterministic elongation, with *L* → ∞ at constant 〈*T*〉, Eq. (D.1) simplifies to

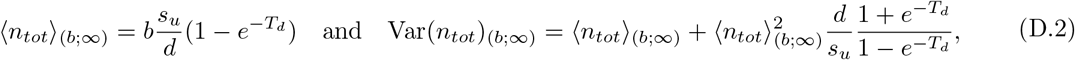

where the subscript (*b*;∞) denotes the bursty limit with infinite *L*. In the limit of zero RNAP detachment, Eq. (D.2) further simplifies to

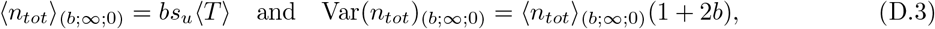

where the subscript (*b*; ∞; 0) denotes the bursty limit, with *L* → ∞ and *d* → 0.

## Moments of total RNAP in the constitutive limit

In the constitutive limit, Eq. (10) simplifies to

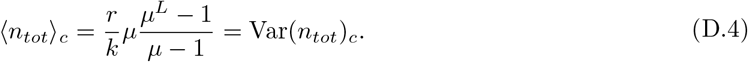

If, furthermore, we take the limit of deterministic elongation, i.e. *L* → ∞ at constant 〈*T*〉, Eq. (D.4) simplifies to

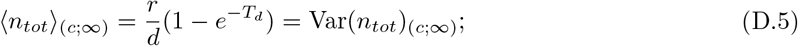

finally, in the limit of zero RNAP detachment, Eq. (D.5) further simplifies to

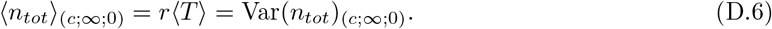

## Moments of mature RNA distribution in the bursty limit

In that limit, the closed-form expressions in Eq. (8) are given by

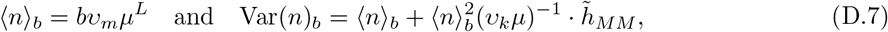

which in the limit of deterministic elongation simplify to

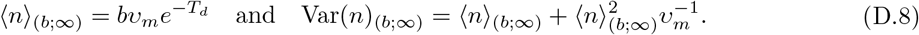

In the limit of zero RNAP detachment, these expressions further simplify to

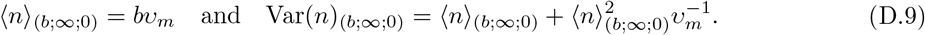

## E Introduction to Geometric Singular Perturbation Theory (GSPT)

We consider a system of first-order autonomous ordinary differential equations in the general (‘standard’) form

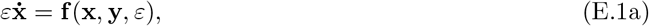

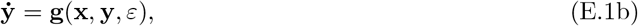

where (**x**, **y**) ∈ ℝ^*m*^ × ℝ^*l*^, with *m, l* ∈ ℕ. Here, 0 < *ɛ* ≪ 1 is a (real) singular perturbation parameter, and the the overdot denotes differentiation with respect to the ‘slow’ time *t*. (Correspondingly, Eq. (E.1) is referred to as the ‘slow’ system.) The variable **x** is referred to as the ‘fast variable’, while **y** is the ‘slow variable’. For simplicity, the functions **f**: ℝ^*m*^ × ℝ^*l*^ × ℝ^+^ → ℝ^*m*^ and **g** : ℝ^*m*^ × ℝ^*l*^ × ℝ^+^ → ℝ^*l*^ are assumed to be 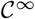-smooth in all their arguments. In the context of our analysis of the characteristic system in Eq. (26), we have the ‘slow system’

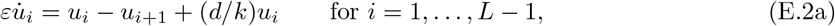

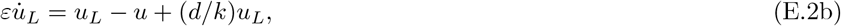

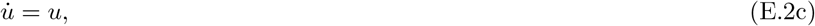

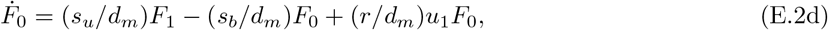

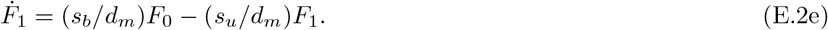

By comparing the system of equations in Eq. (E.2) with the general form in Eq. (E.1), we see that *u*_*i*_ (*i* = 1, …, *L*) are the fast variables, while *u*, *F*_0_, and *F*_1_ are slow. Correspondingly, we have *m* = *L* and *l* = 3 in the above notation, which implies **f** = (*f*_1_, *f*_2_, …, *f_L_*), with *f*_*i*_ = *f*_*i*_ (*u*_*i*_, *u*_*i*+1_) = *u*_*i*_ − *u*_*i*+1_ + (*d/k*)*u*_*i*_ for *i* = 1, …, *L* − 1, *f*_*L*_ = *f*_*L*_(*u*_*L*_, *u*) = *u*_*L*_ − *u* + (*d/k*)*u*_*L*_, and **g** = (*g*_1_, *g*_2_, *g*_3_)(*u*_1_, *u, F*_*0*_, *F*_1_) = (*u,* (*s*_*u*_/*d*_*m*_)*F*_1_ − (*s*_*b*_/*d*_*m*_)*F*_0_ + (*r/d*_*m*_)*u*_1_*F*_0_, (*s*_*b*_/*d*_*m*_)*F*_0_ − (*s*_*u*_/*d*_*m*_)*F*_1_).

Now, we introduce a new ‘fast’ time *τ* = *t/ɛ*, which we substitute into Eq. (E.1) to find the ‘fast system’

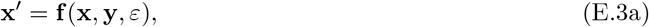

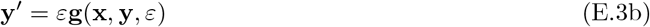

corresponding to Eq. (E.1); here, the prime denotes the derivative with respect to *τ*. Hence, rewriting Eq. (E.2) in the fast formulation, we find

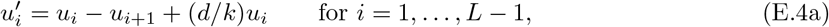

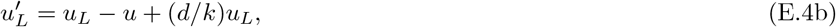

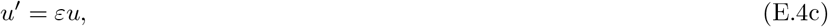

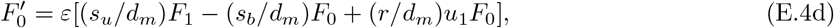

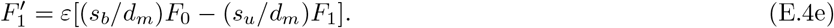

For positive *ɛ*, the systems in Eqs. (E.1) and (E.3) – and, correspondingly, the systems in Eqs. (E.2) and (E.4) – are equivalent; however, in the singular limit of *ɛ* → 0, we obtain two different systems: setting *ɛ* = 0 in Eq. (E.1), we have the ‘reduced problem’

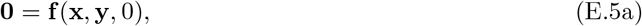

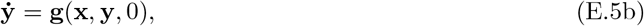

while we obtain the ‘layer problem’

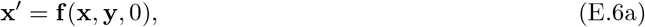

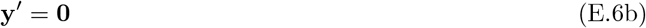

for *ɛ* = 0 in Eq. (E.3). The ‘reduced problem’ for the system in Eq. (E.2) implies that the flow of (*u, F*_0_, *F*_1_) is constrained to lie on the (*l* = 3)-dimensional ‘critical manifold’ 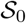 that is defined by **f** = **0**:

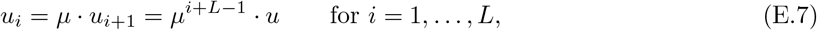

where *u*_*L*+1_ ≡ *u* and (*F*_0_, *F*_1_) are assumed to vary in an appropriately chosen subset of ℝ^2^.

From the ‘layer problem’ of the system in Eq. (E.2), we conclude that **y** = (*u, F*_0_, *F*_1_) is a parameter which parametrizes the (*m* = *L*)-dimensional flow of *u*′_*i*_ = *f*_*i*_ (*i* = 1, …, *L*), the equilibria of which are located on 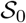.

The Jacobian matrix **D**_x_**f**(**x**, **y**, 0) of the ‘layer problem’ corresponding to Eq. (E.4) about 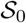 has the eigenvalues

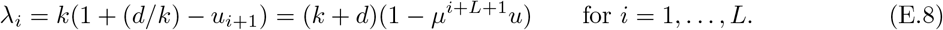

Since our definition of the generating function *F* (*z, τ*) in Section 3.2 assumed *z* ∈ [−1, 1], we may restrict to *u* ∈ [−2, 0] which, by Eq. (E.8), implies that *λ*_*i*_ > 0. Hence, the critical manifold 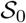 is ‘normally hyperbolic’ – and, in fact, normally repelling – with an (*m* + *l* = *L* + 3)-dimensional unstable manifold 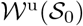.

The geometric singular perturbation theory due to Fenichel [26] thus implies that 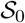 will persist, for *ɛ* positive and sufficiently small, as a slow manifold’ 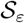 that is (locally) invariant, smooth, and 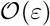-close to 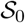. (As the unstable manifold 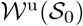 equals the entire phase space of Eq. (E.2), it trivially persists as the unstable manifold 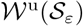 for 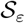.) In particular, as 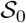 is repelling in forward time, it follows that the inverse characteristic transformation corresponding to Eq. (26) is well-defined in backward time; details can be found in [36, 39].

## F Variance of fluctuating total fluorescent signal

By definition, the variance of the total fluorescent signal is given by the sum over all elements Cov(*r*_*i*_, *r*_*j*_) for *i, j* = 1, …, *L*, where *r*_*i*_ = (*v/L*)*in*_*i*_; the corresponding definitions can be found in Section 4 of the main text. Hence, we have that

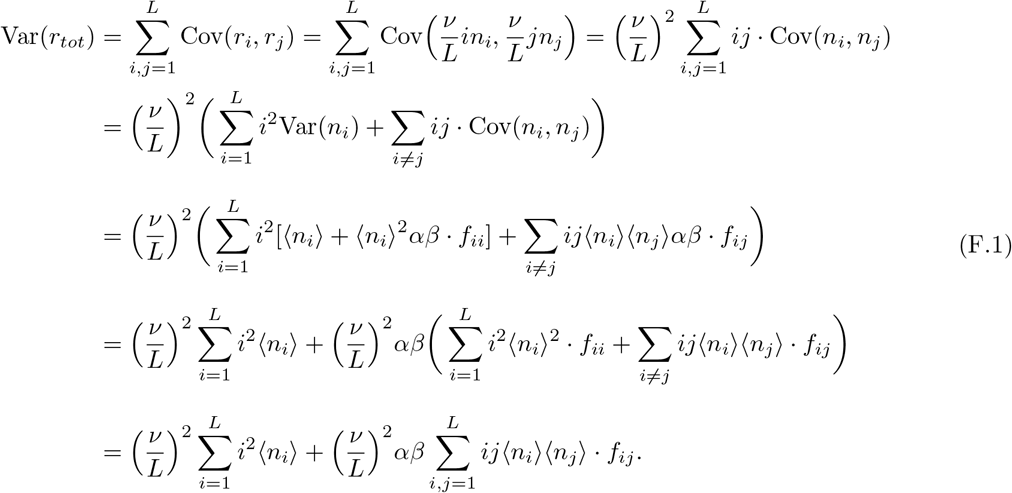

Substituting the expressions for the means 〈*n*_*i*_〉 from Eq. (2b) into Eq. (F.1), we obtain

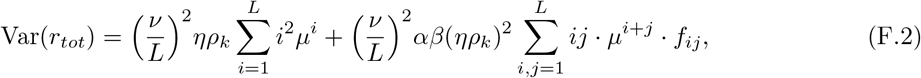

which is the expression stated in Eq. (35).

## G Moments of fluctuations in total fluorescent signal in various limits

## Deterministic elongation

Substituting *k* ↦ (*L*/〈*T*〉 − *d*) and taking the long-gene limit of *L* → ∞ in Eq. (35), we obtain the simplified expressions

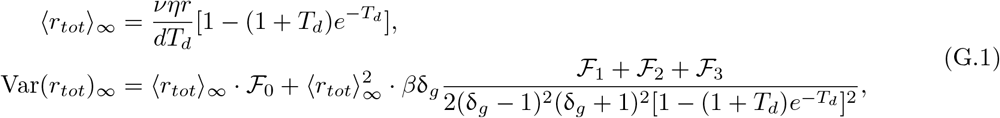

where

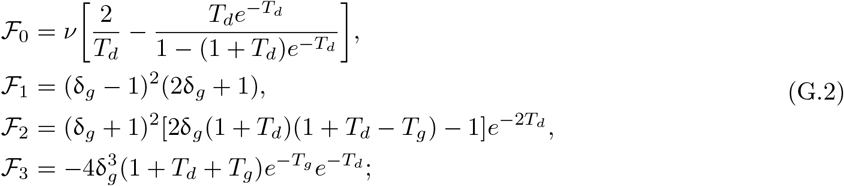

the expression for the variance in Eq. (G.1) is found via the same method as is used in Lemma C.1 of Appendix C. When there is no detachment of RNAP from the gene, i.e. when *d* = 0, Eq. (G.1) simplifies to

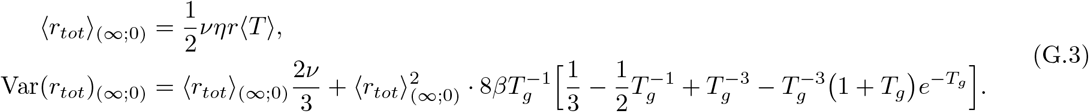

## Bursty limit

In the limit when the rates *s*_*b*_ and *r* are large, the expressions for the mean and variance of the total fluorescent signal given in Eq. (35) become

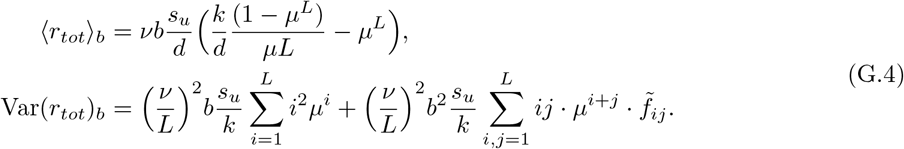

## Constitutive limit

When the gene spends most of its time in the active state, Eq. (35) simplifies to

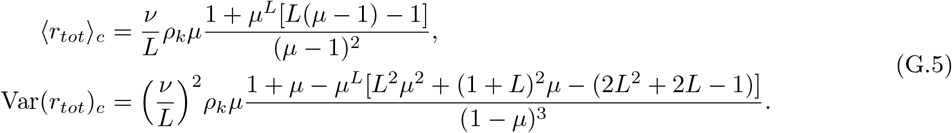

## Bursty expression with deterministic elongation

In that case, Eq. (G.4) simplifies to

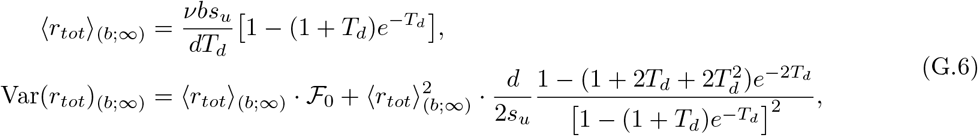

where 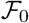 is given by Eq. (G.2). In the special case of zero premature RNAP detachment from the gene (*d* → 0), Eq. (G.6) can be further simplified to

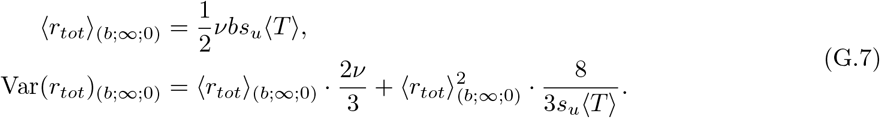

## Constitutive expression with deterministic elongation

In that case, Eq. (G.5) simplifies to

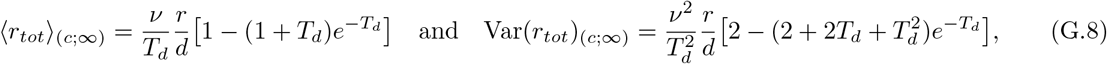

which reduces to

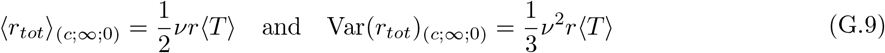

for the special case of zero RNAP detachment from the gene.

## H Extended model with RNAP pausing

*Proof of Proposition 3.* The new pausing model presented in Fig. 9 can be conveniently described by 2*L* + 2 species interacting via an effective set of 5*L* + 4 reactions. The vector 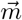 of the number of molecules of the respective species is given by 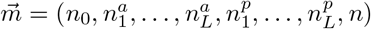; in the table below, we summarize the respective positions of each entry in 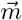, as well as the definition of the rate functions *f*_*j*_, for *j* = 1, …, 5*L* + 4.

| Species | Molecule numbers | Position (in $\vec{m}$) |
| --- | --- | --- |
| $G_{\text{on}}$ | $n_0$ | 1 |
| $P_i, \quad i \in \{1, \dots, L\}$ | $n_i^a$ | $i + 1$ |
| $\bar{P}_i, \quad i \in \{1, \dots, L\}$ | $n_i^p$ | $i + L + 1$ |
| $M$ | $n$ | $2L + 2$ |

| Reaction | Rate function $f_j$ |
| --- | --- |
| $G_{\text{on}} \xrightarrow{s_b} G_{\text{off}}$ | $f_1 = s_b \langle n_0 \rangle$ |
| $G_{\text{off}} \xrightarrow{s_u} G_{\text{on}}$ | $f_2 = s_u (1 - \langle n_0 \rangle)$ |
| $G_{\text{on}} \xrightarrow{r} G_{\text{on}} + P_1$ | $f_3 = r \langle n_0 \rangle$ |
| $P_i \xrightarrow{k} P_{i+1}, \quad i \in \{1, \dots, L-1\}$ | $f_{i+3} = k \langle n_i^a \rangle$ |
| $P_L \xrightarrow{k} M$ | $f_{L+3} = k \langle n_L^a \rangle$ |
| $P_i \xrightarrow{d_a} \emptyset, \quad i \in \{1, \dots, L\}$ | $f_{i+L+3} = d_a \langle n_i^a \rangle$ |
| $P_i \xrightarrow{r_p} \bar{P}_i, \quad i \in \{1, \dots, L\}$ | $f_{i+2L+3} = r_p \langle n_i^a \rangle$ |
| $\bar{P}_i \xrightarrow{r_a} P_i, \quad i \in \{1, \dots, L\}$ | $f_{i+3L+3} = r_a \langle n_i^p \rangle$ |
| $\bar{P}_i \xrightarrow{d_p} \emptyset, \quad i \in \{1, \dots, L\}$ | $f_{i+4L+3} = d_p \langle n_i^p \rangle$ |
| $M \xrightarrow{d_m} \emptyset$ | $f_{5L+4} = d_m \langle n \rangle$ |

Note that we do not consider $G_{\text{off}}$ as an independent species, as a conservation law implies $\langle G_{\text{off}} \rangle = 1 - \langle n_0 \rangle$ . Given the ordering of species and reactions as described in above tables, we can define the $(2L+2) \times (5L+4)$ -dimensional stoichiometry matrix $\mathbf{S}$ , with non-zero elements given by

Note that we do not consider *G*_off_ as an independent species, as a conservation law implies 〈*G*_off_〉 = 1 − 〈*n*_0_〉. Given the ordering of species and reactions as described in above tables, we can define the (2*L* + 2) × (5*L* + 4)-dimensional stoichiometry matrix **S**, with non-zero elements given by

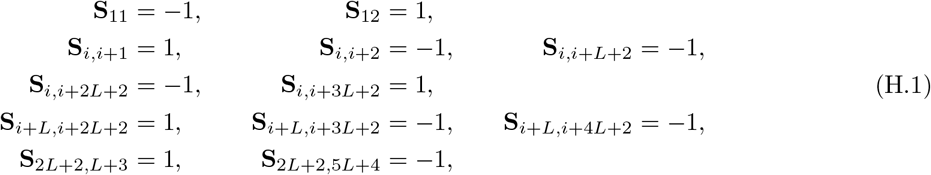

where *i* = 2, …, *L* + 1. From the associated CME, it can be shown via the moment equations that the time-evolution of the vector 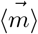 of mean molecule numbers in a system of reactions with propensities that are linear in the number of molecules is determined by 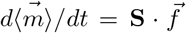. Given the form of the stoichiometric matrix **S** and of the rate functions *f*_*j*_, it follows that the mean numbers of molecules of active gene, active and paused RNAP, and mature RNA in steady-state can be obtained by solving the following system of 2*L* + 2 algebraic equations:

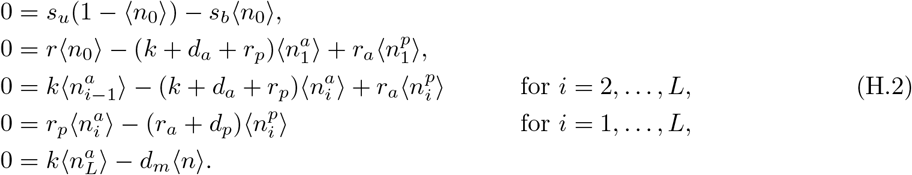

Here, we recall the definition of the following parameters from the main text: *η* = *s*_*u*_*τ*_*g*_, where *τ*_*g*_ = 1/(*s*_*u*_ + *s*_*b*_) is the gene switching timescale, *ρ*_*k*_ = *r/k*, and *ρ* = *r/d_m_*. Also, we define several new parameters: *σ* = *r*_*p*_/*r*_*a*_ as the ratio of the pausing and activation rates; 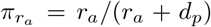, which is the probability of RNAP switching to the active state; 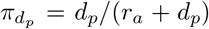, which is the probability of premature termination from the paused RNAP state; 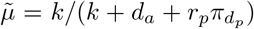; and 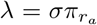. It follows that the solution of Eq. (H.2) can be written as

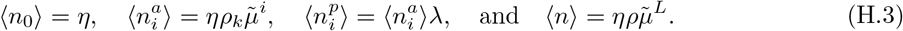

*Proof of Proposition 4.* In order to solve the Lyapunov equation **J** · **C** + **C** · **J**^*T*^ + **D** = **0** for the symmetric elements **C**_*ij*_ = **C**_*ji*_ of the (2*L* + 2) × (2*L* + 2)-dimensional covariance matrix **C**, we will follow the same approach as in Appendix B. First, we define the (2*L* + 2) × (2*L* + 2)-dimensional Jacobian and diffusion matrices for our system. The Jacobian matrix **J** has the following non-zero elements,

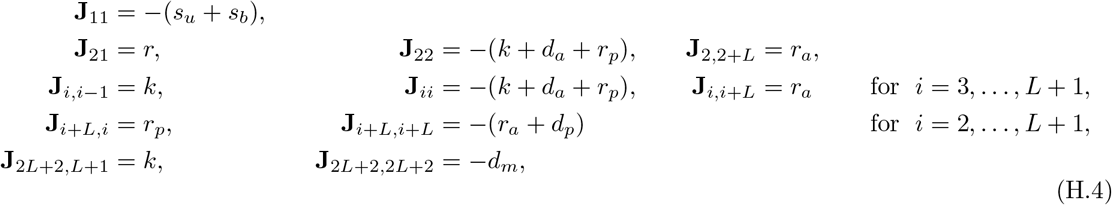

while the non-zero elements of the symmetric diffusion matrix **D** are given by

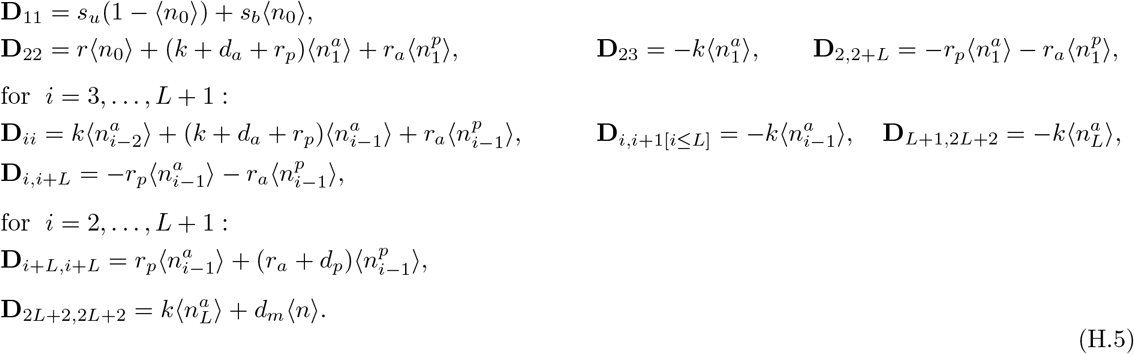

Next, using the definition of **J** and **D** from Eqs. (H.4) and(H.5), respectively, we solve the Lyapunov equation. Here, we note that we are only interested in expressions for the covariances of fluctuations in active and paused RNAP, but not of mature RNA fluctuations; hence, we require closed-form expressions for the elements **C**_*ij*_ with *i, j* ≠ 2*L*+2, which we derive by following the same procedure as in Appendix B.

Now, we recall that *β* = *s*_*b*_/*s*_*u*_ is the ratio of gene deactivation and activation rates, while *τ*_*p*_ = 1/(*k* + *d*_*a*_) is the typical time that an actively moving RNAP spends on a gene segment. Additionally, let 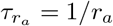 be the timescale of RNAP activation from the paused state, let 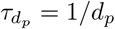 be the timescale of premature termination of paused RNAP, and let *τ*_*pp*_ = 1/(*r*_*a*_ + *d*_*p*_) be the typical time spent in the paused state. Finally, we define the following new parameters: 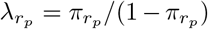, where 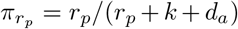 is the probability of actively moving RNAP switching to the paused state, as well as

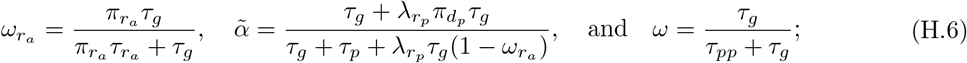

then, closed-form expressions for the covariances of the active gene with itself and the remaining species are given by

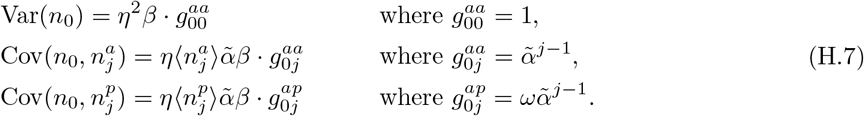

Similarly, closed-form expressions for the covariances between all RNAP species read

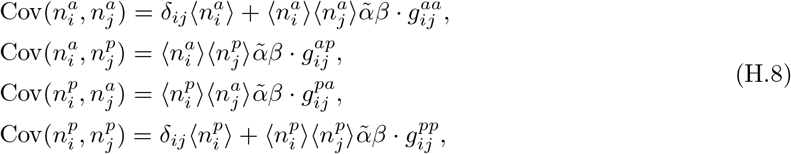

where the functions 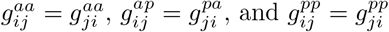 satisfy the following recurrence relations:

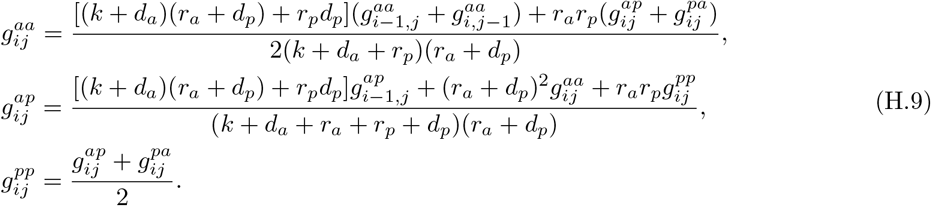

Now, we assume that the elongation rate is faster than the rates of RNAP pausing, activation, and premature termination, i.e. that *k* ≫ *r*_*a*_, *r*_*p*_, *d*_*a*_, *d*_*p*_ in Eq. (H.9). Taking the limit of *k* → ∞, we find that the expressions in Eqs. (H.7) and (H.8) remain unchanged, while Eq. (H.9) simplifies to

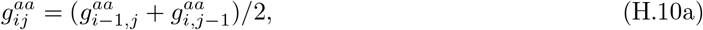

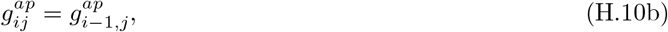

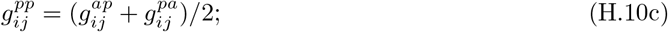

in particular, to leading order in 1/*k*, the functions 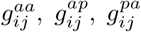, and 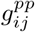 hence do not depend on *k*. Eq. (H.10a) defines a recurrence relation for the symmetric function 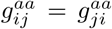 with initial conditions 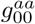 and 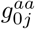 from Eq. (H.7). Using the same mathematical technique as in Lemma B.5, we find that the solution for the function 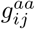 is given by 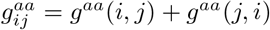, where

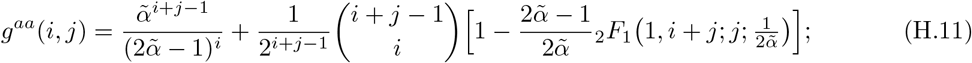

Eq. (H.10b) is a recurrence relation for the function 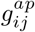 with initial conditions 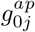 from Eq. (H.7); the corresponding solution is then given by 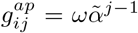. Finally, the solution of the recurrence relation in Eq. (H.10c) for 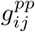 is given by 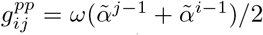. In sum, the leading-order asymptotics (in 1/*k*) of the covariances between the various RNAP species for *k* large is hence given by Eq. (H.8), with 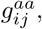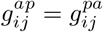, and 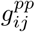 as stated above.

## Asymptotics of variance of total RNAP distribution

The variance of the total RNAP distribution for the pausing model is given by:

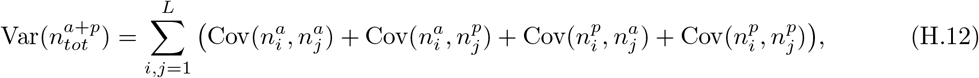

where the expressions for the corresponding covariances are given in Eq. (39). In order to simplify the above expression, we consider each term on the right-hand side in Eq. (H.12) separately, as follows:

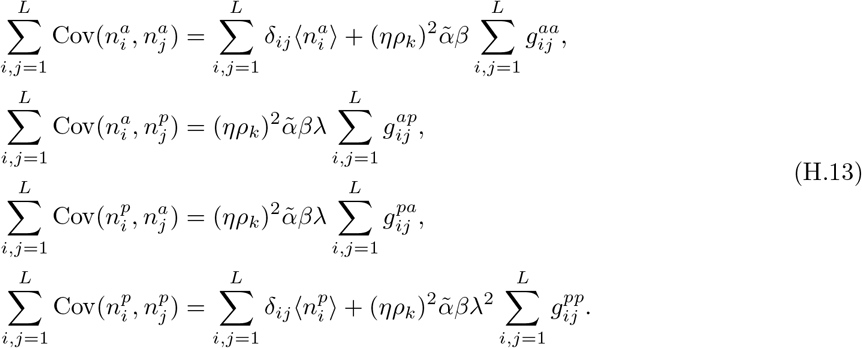

Since 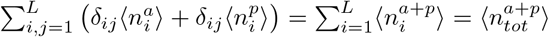, Eq. (H.12) becomes

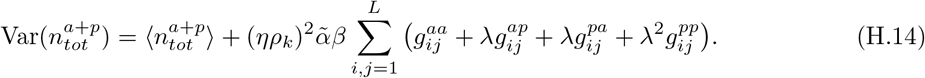

By using the expressions for the functions 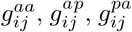, and 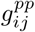 from Eq. (39), we conclude that Eq. (H.14) further simplifies to

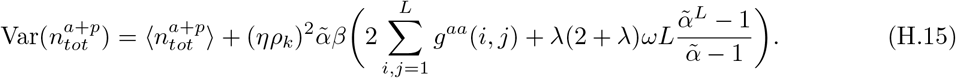

## I Approximation of mature RNA distribution in extended model

Similarly to Section 3.2, we apply geometric singular perturbation theory (GSPT) to formally derive the distribution of mature RNA for the extended pausing model. As was done there, we define 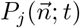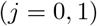 as the probability of the state 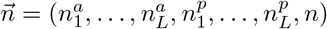 at time *t* while the gene is either active (0) or inactive (1); then, the time evolution of these probabilities can be described by a system of coupled CMEs:

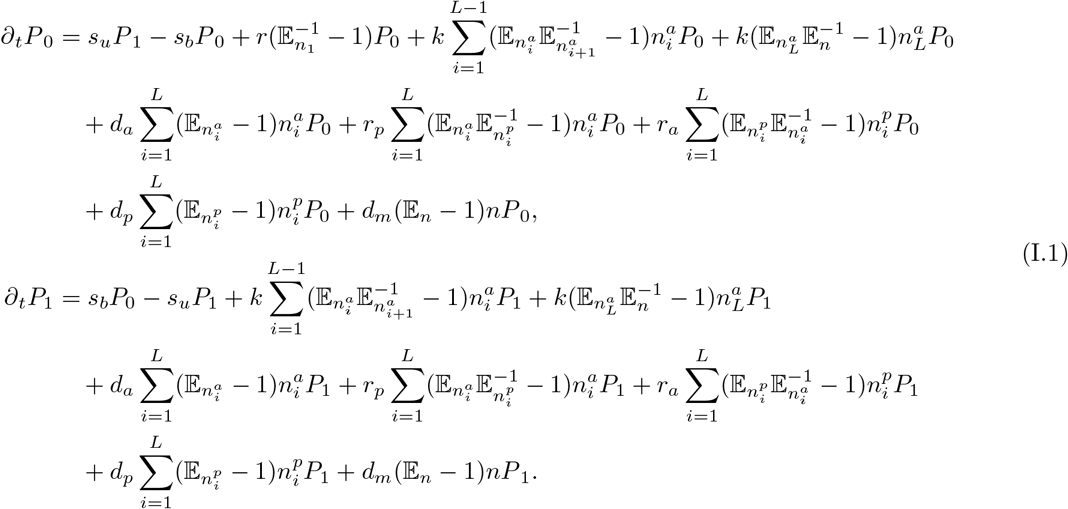

In order to find analytical expressions for the propagator probabilities 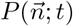 which satisfy the system of CMEs in Eq. (I.1), we define the probability-generating functions 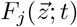, where 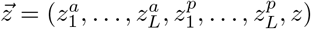 is a vector of variables corresponding to the state 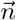. Given the equations for 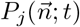 from Eq. (I.1), we obtain the following system of PDEs for the corresponding generating functions 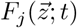:

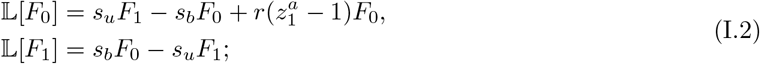

here,

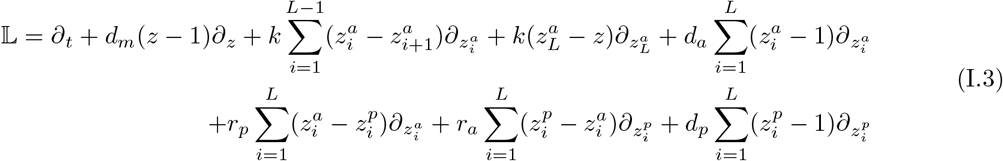

is a differential operator acting on the functions *F*_0_ and *F*_1_. Eq. (I.2) represents a system of coupled, linear, first-order PDEs. Now, we introduce new variables 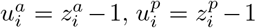, and *u* = *z* − 1; we also rescale all rates and the time variable with the degradation rate *d*_*m*_ of mature RNA. Next, we apply the method of characteristics, with *s* being the characteristic variable. The first characteristic equation will give us *d*_*m*_(*dt/ds*) = 1, with solution *s* ≡ *d*_*m*_*t*; hence, we can use the variable *t*′ = *d*_*m*_*t* as the independent characteristic variable and thus convert the system of PDEs in Eq. (I.2) into a characteristic system of ODEs:

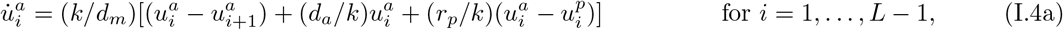

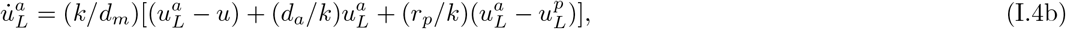

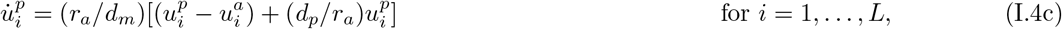

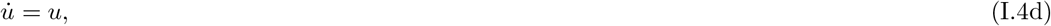

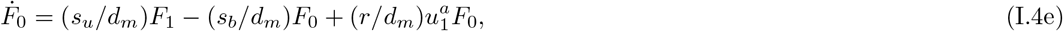

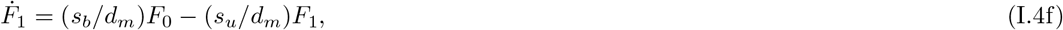

where the overdot denotes differentiation with respect to *t*. Here, we assume that *k*/*d*_*m*_ ≫ 1 and *r*_*a*_/*d*_*m*_ ≫ 1; hence, we define *ɛ* = *d*_*m*_/*k* as the singular perturbation parameter, and we write *d*_*m*_/*r*_*a*_ = *ɛδ*, where 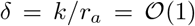 by assumption. Since 0 < *ɛ* ≪ 1 is small, we can apply GSPT in order to separate the system in Eq. (I.4) into fast and slow dynamics, which will allow us to find an asymptotic approximation for *F*_0_ and *F*_1_ in steady-state. With the above definitions, the governing equations for 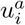 and 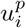 in the ‘slow system’ in Eqs. (I.4a) through (I.4c) become

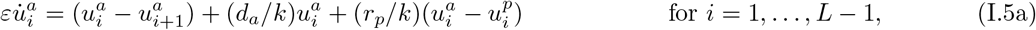

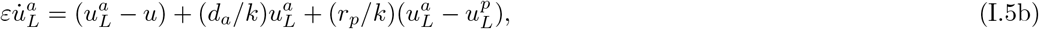

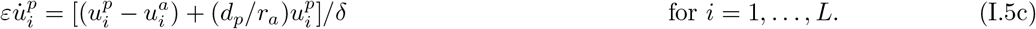

It follows that 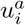 and 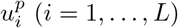 are the fast variables in our system, while *u*, *F*_0_, and *F*_1_ are the slow ones; see Appendix E. Setting *ɛ* = 0 and solving the system in Eq. (I.5), we find 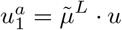, where 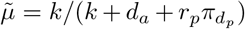 has previously been defined in Prop. 3. Now, given Eq. (I.4d), we apply the chain rule, *dt*′ ≡ *du* · *u*, to rewrite Eqs. (I.4e) and (I.4f) as:

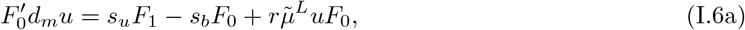

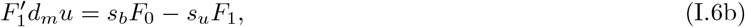

where the prime now denotes differentiation with respect to *u*. The system in Eq. (I.6) is the same as that in Eq. (28), with the substitution 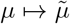; hence, following the same derivation as in Section 3.2, we conclude that the steady-state analytical expression for the probability distribution of mature RNA is given by

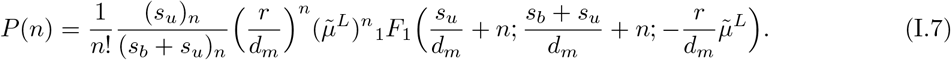

